# Understanding visual attention with RAGNAROC: A Reflexive Attention Gradient through Neural AttRactOr Competition

**DOI:** 10.1101/406124

**Authors:** Brad Wyble, Chloe Callahan-Flintoft, Hui Chen, Toma Marinov, Aakash Sarkar, Howard Bowman

## Abstract

A quintessential challenge for any perceptual system is the need to focus on task-relevant information without being blindsided by unexpected, yet important information. The human visual system incorporates several solutions to this challenge, one of which is a reflexive covert attention system that is rapidly responsive to both the physical salience and the task-relevance of new information. This paper presents a model that simulates behavioral and neural correlates of reflexive attention as the product of brief neural attractor states that are formed across the visual hierarchy when attention is engaged. Such attractors emerge from an attentional gradient distributed over a population of topographically organized neurons and serve to focus processing at one or more locations in the visual field, while inhibiting the processing of lower priority information. The model moves towards a resolution of key debates about the nature of reflexive attention, such as whether it is parallel or serial, and whether suppression effects are distributed in a spatial surround, or selectively at the location of distractors. Most importantly, the model develops a framework for understanding the neural mechanisms of visual attention as a spatiotopic decision process within a hierarchy and links them to observable correlates such as accuracy, reaction time, and the N2pc and P_D_ components of the EEG. This last contribution is the most crucial for repairing the disconnect that exists between our understanding of behavioral and neural correlates of attention.

## 1 Introduction

A quintessential challenge for any perceptual system is the need to focus on task-relevant information without being blindsided by unexpected information that is also important. For example, a driver must be able to stop in response to an unexpected obstacle even while searching intensely for a specific landmark. Understanding how perception meets this ubiquitous challenge is crucial for understanding how the brain balances the prioritization of sensory information according to its relevance.

This challenge is matched by a multitude of attentional systems operating across different senses and time scales. For example in vision there are overt and covert forms of spatial attention, and within covert attention, there is a further distinction between a rapid transient/reflexive form of spatial attention and a slower sustained/volitional form (Jonides 1981; Muller & Rabbit 1992; Hopfinger & Mangun 1998; Nakayama & Mackeben 1989). There are also non-spatial forms of attention that allow us to select among spatially overlapping visual inputs (Neisser & Becklen 1975). Decades of research have provided a multitude of data types that define the properties of visual attention, such as accuracy, reaction time and neural correlates such as Event Related Potentials (ERPs). These data have driven the development of many theories, but the great majority of them are linked to specific paradigms (e.g. a model of visual search, or a model of the attentional blink). Such models are a useful starting point, but their focus on tasks makes it difficult to generalize across experimental paradigms, and also makes it easy to inadvertently overfit a theory to a specific kind of finding. Newell (1973) argued that instead of focusing on individual kinds of results as a way to attack or defend a theoretical edifice, we can use a collection of results most productively if we build a comprehensive model that addresses all of them. The approach used here is to build a model that is close to the algorithmic level of implementation (Marr 1982) and that maximizes the number of empirical constraints that can be applied (Love 2015) with a minimum of parameter adjustment.

The model described here, termed RAGNAROC, which is short for ***R****eflexive **A**ttention **G**radient through **N**eural **A**tt**R**act**O**r **C**ompetition*, is intended as a theoretical and computational framework for understanding how the visual system implements a reflexive form of attention.

This model addresses data in different forms (e.g. accuracy, reaction time and EEG), and from a diversity of paradigms with a goal of building a formalized understanding of how the visual system rapidly makes decisions about which stimuli to enhance and suppress. Moreover, these mechanisms will be linked to observable neural correlates such as the N2pc and P_D_ components. The model also provides suggested resolutions for ongoing debates in the literature by showing how one model is able to account for seemingly contradictory patterns of data (e.g. simultaneous attention to two stimuli but also suppression of competing representations). For the reader who is more interested in the conclusions than the model methods, there is a section in the discussion that focuses on the lessons that have been learned through the construction of the model.

### 1.1 Scientific philosophy of this account and intended audience

This paper is written with the perspective of the experimental scientist in mind. Equations will be kept to a minimum, except for the appendix, and figures will be used to explain the model’s dynamics. RAGNAROC was developed according to abductive principles of theory design (Haig 2005) in which existing data are used to abduce a causal explanation. Thus the neural mechanisms proposed here are intended to represent the simplest possible solution to explaining such data that are relevant to the mechanisms of reflexive attention, while adhering to constraints of neural plausibility. The goal of abduction is to distill a likely explanation for an existing set of data, and this explanation can then be tested through further empirical work.

In terms of validation, we consider the problem to exist in the M-open class (Clarke, Clarke & Yu 2013), which is to say that it is impossible to exactly specify the biological system in this context. Nevertheless, abstract neural models such as this one are a powerful way to distill insights and predictions to guide future research. The paper concludes with a set of lessons and predictions that should be of interest to anyone who studies visual cognition..

### 1.2 Model scope

This model is not to be taken as a complete model of visual attention, which would be beyond the scope of any single paper. RAGNAROC does not address, for example, how attentional control affects eye movements (Rao, Zelinsky Hayhoe, & Ballard 2002; Zelinksy 2008) or slower forms of covert attention that are more firmly under volitional control and can be maintained for a prolonged duration (e.g. multiple object tracking Pyslyshyn & Storm 1988). The model includes a mechanism for the enhancement of information processing, but this is intended only as a proxy for more comprehensive explanations that would interface more directly with single-unit data (e.g. Reynolds & Heeger 2009; Beuth & Hamker 2015). Another variety of attentional mechanisms not addressed here are those that track object features rather than spatial locations (e.g. Neisser & Becklen 1975; Blaser, Pylyshyn & Holcombe 2000). Also, the model explains the initial deployment of attention at the onset of a display which presumably plays a role in visual search. However, the model does not address the goal-driven iterative attention processes that occur over a longer time scale and would be required during many visual search tasks.

In terms of anatomy, we describe the reflexive attentional system in terms of a hierarchy of multiple maps that is inspired by work on the macaque posterior cortex and reinforced by the fact that the lateralized EEG components associated with attention discussed here are also primarily posterior in origin. However it is likely that a combination of frontal and subcortical areas are involved in these processes, and it is not our intent to suggest that reflexive attention is exclusively mediated by posterior areas. Moreover, the model is focused on attention effects at approximately the time scale of one fixation, and thus is not directly applicable to tasks that require multiple cycles of attentional engagement.

### 1.3 The complexity of understanding attention

In broad strokes, attention is perhaps best summarized as privileging certain representations at the expense of others and this prioritization takes many forms throughout the nervous system, ranging from internal control signals within the brain all the way down to active sensing by orienting concentrated receptor clusters, such as the finger tips and fovea, toward relevant stimuli. In terms of visual attention, a distinction is often drawn between *voluntary attention*, wherein volitional control mechanisms configure the spatial deployment of attention over an extended period of time and *reflexive attention*, wherein the visual system reacts rapidly to stimulus onsets in order to attend them before the stimulus display changes or the eye moves (Jonides 1981; Muller & Rabbit 1992; Hopfinger & Mangun 1998; Nakayama & Mackeben 1989). The term *reflexive* invokes an analogy with muscle reflexes that are deployed rapidly in response to a stimulus, and without waiting for slower, deliberative processes.

This reflexive form of attention presumably plays a key role in selecting important information for further processing when the eyes are making saccades frequently. Moreover, it is known to be responsive to higher levels of cognitive control, such that goals, expectations and rewards moderate how strongly stimuli can trigger or capture attention (Folk, Remington & Johnston, 1992). What we do not yet understand is how such a rapid form of attention would function at the level of neural mechanisms.

The earliest theories of such attentional effects described how attention can be distinguished into a series of operations including a generalized alerting function, localizing a target, engaging attention with a stimulus, disengaging attention and finally inhibiting that attended location (Posner, Inhoff, Friedrich & Cohen 1987). Later theories elaborated these mechanisms by proposing that attention involves a combination of target enhancement (Eimer 1996), and suppression of distractors (Cepeda, Cave, Bichot & Kim 1998, Gaspelin Leonard & Luck 2015). However, while it seems straightforward to postulate such attentional effects, these operations are non-trivial to implement in a visual system that is distributed across cortical regions. In such a system, it is not immediately obvious how neural representations could be tagged as belonging to a target or distractor. Furthermore, how does the brain implement such a coordinated attentional process across this network of interconnected maps without requiring an exhaustively large number of intra-cortical connections? Limitations on white matter density imply that it is not feasible for all neurons to communicate directly with all other neurons, which makes seemingly straightforward decision-making approaches such as *winner-take-all* (i.e. the strongest representation suppresses all others) impractical.

An additional complication arises when we consider that the attentional system cannot afford to implement a crisp categorical distinction between targets and distractors. No matter how strongly a person is engaged on a task, there must always be a possibility for task-irrelevant information to trigger attention so that the system remains responsive to unexpected dangers.

Thus, it must be the case that all stimuli, whether designated as targets or distractors by the experimental paradigm, are evaluated to some degree. One might be tempted to argue that inattentional blindness experiments (Rock, Linnett, Grant & Mack 1992; Neisser & Becklen 1975; Simons, & Chabris 1999) demonstrate effective suppression of unexpected information. However, many subjects do notice the unexpected stimulus in such experiments. Moreover, the proportion of participants who noticed, for example, the black gorilla in Simons & Chabris (1999), was influenced by the attentional set of the observer. Furthermore, some studies found that the unreportable stimuli in inattentional blindness could influence perception (e.g., Moore & Egeth, 1997).

### 1.4 Behavioral evidence for covert attentional control mechanisms in vision

#### 1.4.1 Reflexive Attention

Reflexive attention is likely to play a role in many visual tasks, and its effects can be observed in paradigms that produce attentional cueing (Posner 1980; Chen & Wyble 2018) attentional capture (Theeuwes 1991, Folk, Remington & Johnston 1992; Yantis 1996) and the early lags of the attentional blink (Shapiro, Raymond & Arnell 1992; Chun & Potter 1995) (Figure 1). In these paradigms, the effect of attention varies according to the nature of the stimuli and the required response. For example, a visual cue increases the accuracy and decreases reaction times for a subsequent target at the cued location, while having the opposite effect for targets at uncued locations. In attentional capture paradigms, a highly salient distractor causes slower and/or less accurate report of a target presented at a different location and enhanced report of a target at the same location as the salient singleton (Folk, et al. 1992). In attentional blink paradigms, when two targets (T1 and T2) are presented sequentially at a Stimulus Onset Asynchrony (SOA) of about 100ms or less, the second target is easy to see but only when the two targets are presented at the same location (Visser Bischof & DiLollo 1999; Wyble & Swan 2015).

**Figure 1.**
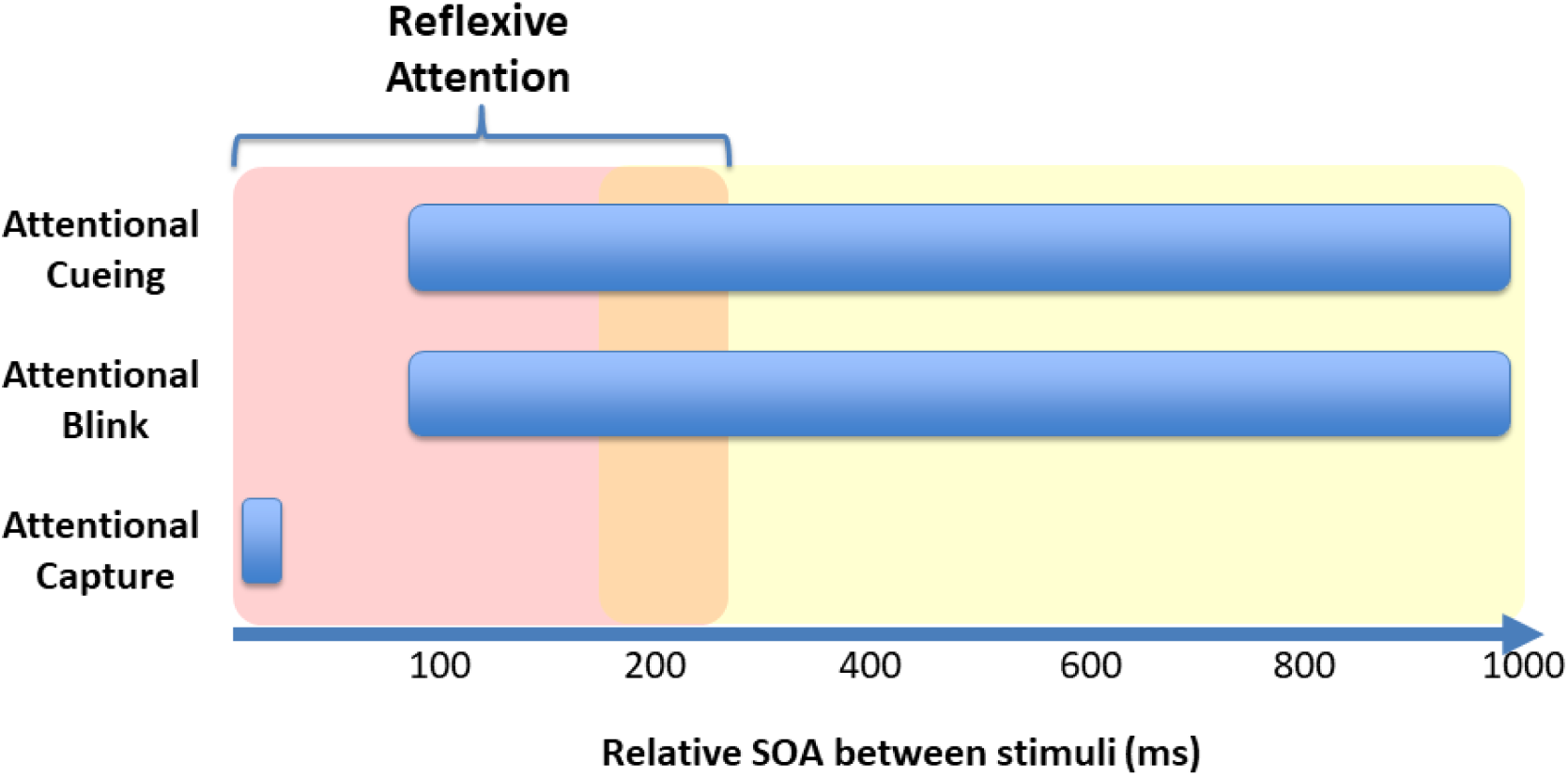
Paradigms that measure attentional effects often present two simultaneous or sequential stimuli and measure the influence of one on the other (e.g. a cue followed by a target, or a T1 followed by a T2). The blue bars indicate the inter-stimulus temporal separations that are most typically studied for three common paradigms. The red portion on the left is the temporal interval over which we consider reflexive attention to play a dominant role in the effect of one stimulus on the other. Attentional effects that involve more volitional forms of processing are dominant at longer synchronies.

#### 1.4.2 Reflexive attention as semi-autonomous control

During normal visual function, the brief duration of eye movements requires a form of attentional control that makes rapid decisions without waiting for confirmation from slower, volitional forms of cognitive control. In this view, reflexive attention is a solution to the demands of the saccadic visual system in that it provides a semi-autonomous decision-making process for selecting information from prioritized locations in the visual field. Reflexive attention is *semi-autonomous* in the sense that the decision process obeys a configuring attentional set that modifies how readily different stimulus attributes will trigger attention. Such attention is strongly driven by salient singletons despite efforts of the subject to ignore those stimuli^1^ (Remington, Johnston, & Yantis 1992) but the likelihood of a stimulus to capture attention is also strongly affected by the similarity between a stimulus and the current goals of the subject (Folk, Remington & Johnston, 1992; Woodman & Luck 1999; Egeth, Leonard & Leber 2011), and to some degree the amount of reward a stimulus has received (Anderson, Laurent & Yantis 2011).

This task-based configuration is even responsive to categorical signifiers such as letters among digits (Wyble, Potter, Bowman 2009; Nako, Wu & Eimer 2014), and superordinate concepts (e.g. “marine animal” Wyble, Folk, Potter 2013). Likewise, neural data from EEG reflect what are thought to be rapid attentional responses to task relevant colors (Eimer 1996); letters/words (Eimer 1996; Tan & Wyble 2015; Nako, Wu & Eimer 2014; Callahan-Flintoft & Wyble 2017) and line drawings (Nako, Wu, Smith & Eimer 2014). Therefore, this system provides a tight coupling between bottom-up (i.e. attention as driven by physical characteristics of the stimulus) and top-down (i.e. attention as driven by expectations, goals and rapid learning) determinants of attentional control. This task-defined specificity coupled with the rapidity of reflexive attention provides a potent way for attention to select task-relevant information even when stimuli are changing rapidly (e.g. Potter 1976; Schneider & Shiffrin 1977).

Despite the fact that reflexive attention can be configured by top down signals, the partial autonomy of this system is evident in the phenomenon of attentional capture, in which attention is deployed to stimuli that appear in locations of the visual field that are known to always be task-irrelevant(Remington, Johnston & Yantis 1992; Theeuwes 1992; Wyble, Folk & Potter 2013; Folk, Leber & Egeth 2002). If reflexive attention were not semi-autonomous, top down control signals would be able to ensure that stimuli presented in locations known to be irrelevant would have no effect on behavior. Note that there are some cases in which top down control settings seem to eliminate salience-based capture (Bacon & Egeth 1994).

Another indication of automaticity comes from Krose & Julesz (1989) who demonstrated that cueing effects were localized to the specific location of a cue in a ring of stimuli, even when subjects were informed that the location of the target would typically be opposite to the location of the cue on the ring (see also Jonides 1981). Thus, expectation induced by both task instructions and experience with the task were unable to eliminate the immediate, reflexive deployment of attention to the specific location of cues at Cue-Target SOAs up to 260ms^2^.

Along similar lines, a finding in electrophysiology by Ansorge, Kiss, Worschech & Eimer (2011) showed that spatial cues that are never in the target’s position will nevertheless generate an N2pc component, with an amplitude that is weighted by top-down feature settings.

A further line of evidence for automaticity is found in a series of experiments in which the cue (a pair of lines) was much larger than the target, and the subject could, in principle, learn how the cue’s properties (e.g. color or shape) determined which part of the cue indicated the likely location of the target (Kristjansson & Nakayama 2003; Kristjansson, Mackeben & Nakayama 2001). It was found that subjects could learn simple relationships, such as that part of a cue (e.g. its left or right half) was more likely to cue a target’s location if that relationship remained consistent across trials. However their attentional allocation was unable to accommodate very simple alteration sequences.

Reflexive attention is also limited in terms of its duration, which is limited, even when it would be advantageous for attention to remain engaged for a longer time period. A good example of this is the transient attention demonstration of Nakayama & Mackeben, (1989) in which, a cue appeared, and stayed on the screen to indicate the location of the target. Even though this cue stayed on the screen and was perfectly predictive of the target location, targets that occurred in the 200ms window after cue onset were reported more accurately than targets appearing at later time points. This effect was replicated by Wilschut, Theeuwes & Olivers (2011) though with a smaller magnitude. This transient effect is not merely an alerting effect since it is spatially selective (Müller & Rabbitt 1989).

#### 1.4.3 Processing enhancement at a cued or target location

Several independent lines of research suggest that deploying reflexive attention enhances the processing of targets in the same location. For example, spatial cueing paradigms find that relative to an uncued condition, a cue will reduce the reaction time to respond to a probe at that location (Eriksen & Yeh 1985) or increase the accuracy of responding to a masked target at that location within about 100ms (Nakayama & Mackeben 1989; Cheal, Lyon & Gottlob 1994; Wyble, Bowman, Potter 2009). The key defining characteristic of the rapid onset of attentional enhancement seems to be that the cue and target appear at the same location, which dovetails with the semi-autonomous nature of reflexive attention. Another case of reflexive attention occurs when two targets are presented in close succession. If they are at the same location and at an SOA of ∼100ms, the second target report is enhanced. Wyble Bowman & Potter (2009). This spatially localized enhancement of processing is also consistent with the finding that lag-1 sparing effects in the attentional blink are strongly linked to spatial congruence between T1 and T2 (Visser Bischof & DiLollo 1999).

#### 1.4.4 Suppression at the location of distractors

Because distractors are, by definition, not explicitly reported or responded to, it has been more difficult to understand how they are affected by attention. One source of information has been to record directly from neurons within the visual system and there are indications in neurophysiology that representations elicited by distractors are suppressed. In single-unit data from monkeys, neurons responsive to a distractor exhibit a sharp reduction in firing rate after about 100 ms when presented alongside a target in the visual field (Chelazzi, Miller, Duncan & Desimone 1993). This finding has been taken as evidence that targets and distractors engage in a competition that is biased towards the target (Desimone & Duncan 1995).

In human behavior, evidence of distractor inhibition in response to a target takes two forms. First, information is suppressed in the surrounding vicinity of a target, as demonstrated when subjects report two targets presented in rapid sequence. These methods reveal an effect termed Localized Attentional Interference (LAI), such that the second target is reported most accurately when in the same position as the first target, much less accurately in the area surrounding the first target (∼3 degrees) and more accurately again at farther separations (Mounts 2000). Bahcall & Kowler (1999) presented a similar finding in which two simultaneous targets were presented at various separations. Cutzu & Tsotsos (2003) also reported a similar finding using a cue and a single target.

In addition to spatial inhibition in the surrounding vicinity of a target, suppression is also centered at the spatial location of distractors. Cepeda, Cave, Bichot & Kim (1998) found that when distractors were presented concurrently with a to-be-reported target, a subsequent probe would be reported more slowly at the location of that distractor, compared to a previously blank location. The implication is that the distractors in the display were suppressed and this suppression carried forward in time to impede the processing of probes presented at the same location. Hickey & Theeuwes (2011) showed that the effect of a distractor that captures attention is greater when spatially proximal to a target, which also implicates a proximity based form of inhibition, centered at the location of a highly-salient distractor. Similarly, Gaspelin, et al. (2015) found that probe letters in a spatial array following or coincident with a search display were harder to report if there had previously been a salient distractor at the location of that letter (although it is crucial to note that this only occurred when participants knew which specific feature to look for; this point will be discussed later). Thus there are two lines of evidence for active inhibition, one locked to the region surrounding a target, and the other centered at the location of distractors. The model presented below will attempt to reconcile these two forms of evidence.

### 1.5 Electrophysiological correlates of visual attention

An important complement to the behavioral evidence of reflexive attention are studies that use Event Related Potential (ERPs) extracted from the EEG, and likewise Event Related Fields (ERFs) from the MEG. ERPs and ERFs provide a measure that is precisely timed to underlying neural events and thus provides crucial information about the relative timing of attentional processes.

#### 1.5.1 ERPs reflecting the current location of spatial attention

When spatial attention has been directed to a specific location prior to the onset of a stimulus, the ERPs evoked by that stimulus will differ according to whether it is inside or outside of the attended location. For example, components elicited by the onset of a visual stimulus such as the N1/P1 complex, are larger in amplitude for a stimulus that appears in an attended location (Mangun 1995; Hillyard & Anllo-Vento 1998) and presumably reflects increased neural activity evoked by stimuli at those locations. Likewise, increased amplitude of the Steady State Visual Evoked Potential (SSVEP) for a flickering stimulus has served as a robust indicator of the location of attention and can last multiple seconds (Müller & Hillyard 2000). These effects indicate that ongoing spatial attention affects the processing of stimuli at the earliest levels of cortical processing. Moreover, they are also useful for demonstrating when shifts of attention have occurred, as in Hopf, Boehler, Luck, Tsotsos, Heinze & Schoenfeld (2006), who demonstrated a neural correlate of the spatial distribution of surround suppression evoked by an attended stimulus (Mounts 2000).

#### 1.5.2 ERPs indicating a change in the spatial distribution of attention

Another class of EEG component is thought to indicate the neural mechanisms involved in the initiation of attention. These potentials, termed the N2pc and the P_D_, occur later in time than the modulations of the N1/P1, which is consistent with the idea that they reflect changes in attention evoked by new stimuli.

##### 1.5.2.1 The N2pc component

The N2pc is a brief negativity elicited by a laterally presented target stimulus, and with an analog that can be recorded for stimuli along the midline (Doro, Bellini, Brigadoi, Eimer & Dell’Acqua 2019). The evoked potential is small in amplitude and occurs on posterior areas of the scalp contralateral to the target, approximately 200-300ms after the onset of the target, and is typically less than 100ms in duration. The original theories proposed by the seminal publications on the N2pc suggested that it reflects either the suppression of distractors (Luck & Hillyard, 1994) or the enhancement of the target (Eimer, 1996). The N2pc has also been referred to as a Posterior Contralateral Negativity or PCN (Töllner, Rangelov & Müller 2012).

Newer findings have provided different perspectives. For example, it has been suggested that the N2pc reflects the process of individuating the target from surrounding stimuli, as its amplitude increases with the number of presented targets, but only when target numerosity is task relevant (Pagano & Mazza 2012). Also, Hickey, Di Lollo, & McDonald, (2009) suggested that when a target is paired with a contralateral distractor, the N2pc to the target is composed of two dissociable components: a negativity evoked by the target (the Nt) and a positivity evoked by the distractor (the P_D_). Since the N2pc is measured as a difference wave between target-contralateral and target-ipsilateral sides of the scalp, the P_D_ would be measured as a negativity relative to the target, thus contributing to the N2pc amplitude. The Nt and P_D_ components were isolated by presenting the distractor or the target, respectively, in the middle of the display, which eliminates their contributions to the ERP and reveals the neural signature evoked by the other stimulus.

Another perspective on the N2pc stems from a finding in which two sequential targets are presented at either the same or different locations on the screen (Tan & Wyble 2015). In the same-location condition, subjects could easily see the second target, however it elicited no additional N2pc beyond the N2pc evoked by the first target. In contrast, when the second target was on the opposite side of the display, a strong second N2pc was evoked by that second target. In terms of behavior, subjects were better at reporting the same-location target, which did not evoke an N2pc, compared to the different-location target which did evoke an N2pc. It was concluded that the N2pc indicates the process of locating a to-be-attended stimulus, rather than enhancement or suppression. This explains the finding that the N2pc was missing for same-location targets, since the second target inherits the attention deployed by the first target, and no additional N2pc is evoked. This effect was subsequently replicated in a series of experiments (Callahan-Flintoft, Chen & Wyble 2018) with the caveat that the second T2 could evoke a very small N2pc, although it was accelerated in time relative to the T1’s N2pc and much smaller in amplitude than that evoked by the T1. This was found to be a consistent with the model of Tan & Wyble (2015) with a small parameter change.

Importantly, this diminished-N2pc phenomenon occurs only when T1 and T2 are presented closely in time. At longer temporal separations (e.g. 600 ms), the T2 elicits a second N2pc, even if subjects have a clear expectation that the target will occur in that location (Callahan-Flintoft, Chen, & Wyble 2018). This finding is crucial because it underscores that reflexive attention is driven by a stimulus, and cannot be maintained for an extended period of time without stimuli to keep attention engaged.

Thus, apart from being related to attention there is little consensus as to the underlying cause of the N2pc. Moreover, a crucial complexity of the N2pc literature is that distractors evoke an N2pc in certain cases (Hickey McDonald & Theeuwes 2006; Burra & Kerzel 2013; Kiss, Grubert, Petersen, & Eimer, 2012; McDonald, Green, Jannati, & Di Lollo, 2012; Liesefeld, Liesefeld, Töllner, & Müller 2017). Such findings highlight the complexity of attentional mechanisms and the difficulty of ascribing unitary functions to neural correlates.

##### 1.5.2.2 The P_D_ Component

Another ERP related to attentional control is the P_D_; a positivity evoked in posterior scalp regions that are contralateral to a distractor (Hickey, McDonald & Theeuwes 2009). The fact that distractors selectively elicit a P_D_ is additional evidence that inhibition is selectively deployed at the location of distractors and reinforces the idea that attention has mechanisms for both enhancing and suppressing information in a spatially selective format.

However, when target search is made extremely easy by re-using the same target-defining feature on each trial and using many repeated trials, salient distractors can be ignored entirely, without producing a P_D_, or an observable behavioral cost (Barras & Kerzel 2016) suggesting that in some cases distractors can be simply ignored rather than suppressed. In other cases salient distractors elicit a P_D_ in the EEG and a minimal cost on the speed of finding the target (Burra & Kerzel 2013) with a concomitant suppression of the distractor’s location as measured by behavioral probes (Gaspelin, Leonard & Luck 2015; Gaspelin & Luck 2018). These results suggest that sometimes the distractor has the potential to interfere, and is inhibited to reduce its influence. Finally, if the search task is made sufficiently difficult by varying the target’s defining feature from trial to trial, then the distractor evokes an N2pc, while also eliciting a strong behavioral capture cost (Burra & Kerzel 2013) suggesting that in such cases there was a consistent failure to inhibit the distractor. This distractor-induced N2pc also supports the suggestion that distractors have an inherent salience which must be inhibited (Sawaki & Luck 2010).

These findings can be summarized as follows. When search is made extremely easy by using many trials and highly prescriptive visual targets, the visual system learns to exclude some kinds of distracting information without reflexive attention. When the task becomes more difficult, distractors are suppressed by spatial inhibition mechanisms, eliciting a P_D_ but no behavioral cost on target response. With a further increase in difficulty by using unpredictable singleton targets, distractors are not as effectively suppressed, allowing them to produce an N2pc and a sizeable behavioral capture effect. Such findings complicate the straightforward attribution of the P_D_ as a correlate of distractor suppression but also underscore the importance of building integrative theories that combine behavioral and neural forms of evidence. As we will argue below, these divergent findings can be explained as a range of outcomes that arise from the competition for attention between putative targets and distractors in a spatially topographic attentional priority map.

## 2. Computational architectures for reflexive attention

Moving to a discussion of how reflexive attentional control might be implemented in the brain, we begin by considering several architectures that could support the ability to selectivity enhance and suppress information acquisition from different locations of a visual display.

### 2.1 Assumptions

This discussion is predicated on several assumptions that are implicit in existing models of attention. An anatomical assumption is that the visual system is hierarchically organized, beginning with low level feature extraction in cortical area V1 that projects to various cortical areas specialized for more specific kinds of information, such as color, and various forms (Van Essen & Maunsell 1983, Kravitz, Saleem, Baker, Ungerleider, & Mishkin, M. 2013; Konkle & Caramazza 2013). These higher-level representations are assumed to maintain the spatial topography of V1 albeit with larger receptive fields (DiCarlo & Maunsell 2003; Silver, Ress & Heeger 2005). Another crucial assumption is that *there is no indicator that definitively determines which stimuli should be attended.* Instead, the attentional system perceives stimuli with varying combinations of intrinsic salience and task relevance. It then decides which stimuli to attend according to the broader goals of the organism, which may sometimes transcend the specific task imposed by the experimenter (i.e. if there was a visible fire in the laboratory, the subject would presumably notice). In other words, the notional distinction between targets and distractors as imposed by any specific task is not the ultimate designation of stimulus priority as far as the visual system is concerned. The system must decide what will be attended and what will be suppressed on each trial and this decision is not pre-ordained by other systems, except perhaps in cases where the visual search is highly prescriptive and repeated many times (Theeuwes 2012; Burra & Kerzel 2013)

A final crucial assumption is that *there are no a-priori labels as to which neurons are processing to-be-attended vs to-be-ignored stimuli.* When a decision has been made to attend to a stimulus, there must exist an efficient means to rapidly distribute the consequence of that decision across a diverse set of cortical areas. For example, a given neuron in early visual cortex may be firing in response to a stimulus that downstream areas of the visual system have determined should be attended, but how is credit assigned back to that neuron?

Given these assumptions, a candidate model of reflexive attention must include mechanisms for making rapid decisions about where to attend, and also mechanisms that rapidly implement that decision by routing information between different portions of the visual system.

### 2.2 Four potential architectures

It is helpful to understand the advantages and drawbacks of various architectures by which attentional decisions could be communicated in a hierarchically organized visual system. This section outlines four possibilities with a particular emphasis on understanding how they scale up to a brain-sized implementation with many levels of processing and many areas within each level).

#### 2.2.1 Local Attentional Control

The simplest method of reflexive attention is implemented at the local circuit level. In such a model, stimuli are processed separately within different maps (Figure 2a). Representations of each stimulus compete within these cortical areas, and one or more winners of that local competition would be attended. While simple, this architecture has difficulty explaining how stimuli of different kinds can affect one another. For example, attentional capture by a color singleton affects processing of a shape singleton target (e.g. Theeuwes 1991) which requires that decision consequences propagate between maps selective for different kinds of information.

**Figure 2.**
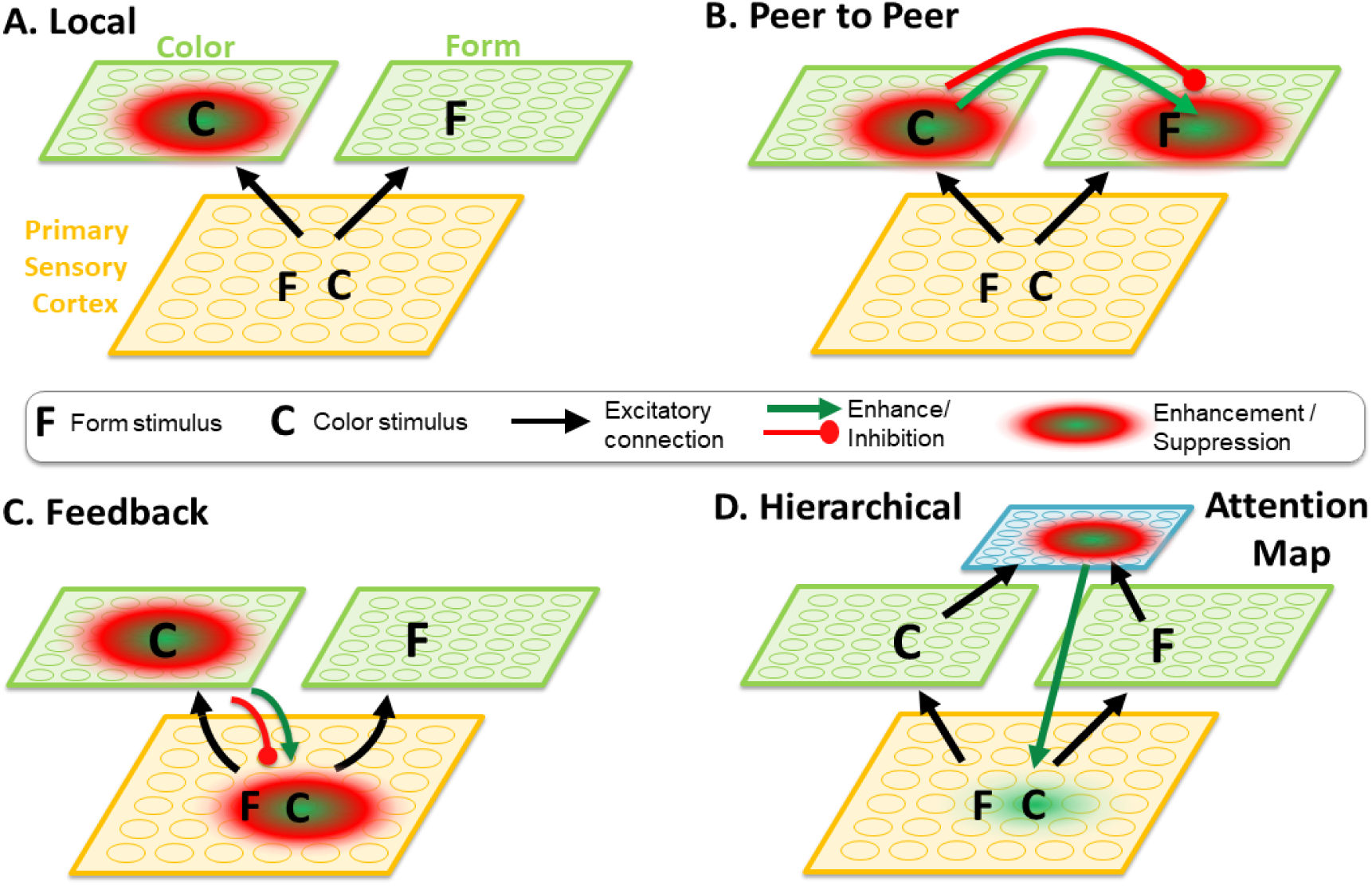
Illustration of four architectures for mediating the competition between two stimuli for which the most salient attributes are processed in different maps (e.g. color singleton vs a form singleton). The illustrations indicate how attentional enhancement and suppression effects elicited by a highly salient color singleton can propagate between areas.

#### 2.2.2 Peer to Peer attentional control

Figure 2b illustrates a model in which any map that resolves a competition between stimuli projects enhancement and suppression to other maps by exploiting the spatiotopic correspondence between feature maps, thus ensuring that the correct locations are excited or inhibited across maps. The disadvantage of this architecture is that it would require an enormous number of intracortical projections at brain scale. Each feature map within the visual system must send dense projections to every other map so that targets in one map can enhance or suppress representations in all other maps. Thus, the number of intra-areal connections grows as M*N^2^, where M is the number of neurons within each cortical area and N is the number of areas. However it has been estimated that only about 30% of the total proportion of possible intra-areal connections exists within the macaque visual system (Felleman & Van Essen 1991) which argues against strong peer-to-peer attentional control.

#### 2.2.3 Feedback Attentional Control

The third architecture is more efficient in terms of intracortical projection (Figure 2c) because it exploits the hierarchical nature of the visual system. Once a stimulus has won a local competition in any map, it projects a combination of enhancement and inhibition back down to the earliest levels of processing in the visual hierarchy (i.e. perhaps V1 or even LGN). These effects then propagate forward to the descendent visual processing areas.

This approach requires fewer inter-areal connections than the peer-to-peer model, growing linearly with the number of feature maps. The selective tuning model of Tsotsos (2011) et al provides a thorough formalization of such a system. The sSoTS model of Mavritsaki, Heinke, Allen, Deco & Humphreys (2011) provides an even more direct example of how stimuli can compete across maps with reference to a master location map in order to determine the most appropriate location to deploy attention.

The disadvantage of this approach is that resolving a competition between two stimuli represented in distinct feature maps requires iterations of feedforward and feedback processing through the hierarchy since the higher level maps do not directly communicate with one another. Furthermore the suppressed information is cut off at the earliest level, which precludes it from analysis by higher levels of the visual system. This makes it difficult for deeper levels of meaning to be computed from stimuli that are not attended.

#### 2.2.4 Inhibition at a superordinate map in a hierarchy

The final architecture that we consider, and the one that is used in RAGNAROC and shared by other algorithms such as SAIM (Heinke & Humphreys 2005; Itti, Koch & Niebur 1998), confines the competition to a single cortical area: an attention map that is hierarchically superordinate to the spatiotopic maps that comprise the ventral visual system (Figure 2d)^3^. The attention map provides a compact method to make rapid decisions about where to deploy attention in the visual field, since it accumulates information about the priority of different stimuli from many subordinate topographic maps into a single brain region. See Rougier (2005) for a similar argument about the advantages of keeping computations local.

Once a spatial region within the attention map has been sufficiently activated by input from subordinate maps, those neurons enhance the processing of information at the corresponding region of the earliest processing level, and that enhancement then carries forward through the ventral hierarchy. In this framework, there is no direct suppression of information in the subordinate layers. Instead, suppression is achieved indirectly by reducing the availability of attention at particular locations in the visual field. Thus, attention is represented as a gradient field of activation levels distributed across the spatial extent of the visual field (LaBerge & Brown 1989, Cheal Lyon & Gottlob 1994). Changes in these activation levels provides a convenient way to throttle the processing of information through all of the feature maps that are descended from the early visual area with a relative minimum of intracortical projections. This approach mitigates the disadvantages of the preceding architectures as follows. Attentional decisions can be made rapidly even between stimuli with distinct representations, since the competition occurs within a single map. Also, suppressing attentional priority, rather than representations in the subordinate layers, preserves the information at the earliest levels of processing, which allows a stimulus in an unattended region the chance to make contact with deeper levels of processing should it be required (i.e. no information is lost).

## 3. RAGNAROC specification

### 3.1 Inspiration from existing models

There is a substantial literature on computational models of attention that collectively addresses a broad set of mechanisms and processes. RAGNAROC is informed by many of these models.

Starting from the very earliest cueing paradigms, there is the idea that attention has a number of stages, and even independent systems for alerting and spatial attention (Posner et al. 1987).

RAGNAROC is inspired by this idea in several respects, firstly that attention involves a number of different mechanisms, and secondly that spatial attention must be localized before it can be engaged. The brief “lock-on” attractor state that emerges from the dynamics of RAGNAROC (more on this below) are akin to LOCALIZE and ENGAGE portions of the Posner et al. model. The DISENGAGE operation also has an analog in the inhibitory Interneurons (more on this below) that help attention to re-allocate to new stimuli.

Another of the formative models for our approach is the Theory of Visual Attention (TVA; Bundesen 1990; Bundesen, Habekost, & Kyllingsbæk, 2011), which provides a mathematical formulation for how goals adjust the pertinence of certain kinds of information (i.e. when one is looking for red digits, the pertinence of red is upweighted to increase the rate at which stimuli with that color are processed). This pertinence weighting applies across the entire visual field, which explains how stimuli are able to capture attention when they match top-down settings despite being located in a to-be-ignored location. TVA also formalizes the understanding of stimulus-driven attention as a decision making process. In terms of implementing a hierarchical architecture for attention, the Koch & Ulman (1985) and Itti, Koch & Niebur (1998) models of salience were crucial for establishing the utility of a shared salience map, which accumulates information from subordinate layers of processing and allows them to compete in a compact neural field. Li (2002) helped to establish the idea of salience being a product primarily of anatomically early levels of processing. Zehetleitner, Koch, Goschy & Müller (2013) elaborated the circuitry of competition at the top of this hierarchy, to provide an illustration of how attention decisions can be considered a race between competing selection operations.

In terms of implementing selection, the Selective Tuning (ST) model of Tsotsos (1995) illustrates how recurrent signals, propagating backwards through the visual hierarchy could implement the selection process at the earliest levels of the hierarchy. However, another class of models has been even more influential in highlighting the importance of recurrence in iteratively shaping the spatial profile of attention. One of the clearest examples of this process is SAIM (Heinke & Humphreys 2003; Heinke & Backhaus 2011) in which a pool of selection neurons interacts with incoming information to create a spatially localized selection and routing of information to a different group of neurons that represent the focus of attention. In SAIM, the selection process is an emergent property of the shared topographic connectivity between the selection system and the retinal input. Input of a given stimulus shape at a given location creates a transient increase in energy that can drive a gradient descent process to ultimately select the stimulus and select the most likely template. This core mechanism of SAIM has recently been shown to approximate the minimization of variational free energy in a Bayesian formulation (Abadi, Yahya, Amini, Friston & Heinke 2018).

In SAIM, a series of top-down connections provides an additional form of resonance that selects for coherent stimuli that match the visual search template. This idea is also present in adaptive resonance models by Grossberg and colleagues, and in particular the attentional shroud model of (Fazl, Grossberg & Mingolla 2009), which describes a process to delineate the boundaries for the purpose of learning.

The idea of reflexive attention, as a brief burst of enhancement to increase processing at a particular moment in time was simulated in the STST (Bowman & Wyble 2007), eSTST (Wyble, Bowman & Nieuwenstein 2009); and Boost Bounce (Olivers & Meeter 2008) models. Those models, especially STST and eSTST, focused more on the time course of encoding of information into memory, whereas RAGNAROC could be conceived as a spatial attention front-end to such models, replacing the simpler “blaster” mechanism that they employed. There has also been work on exploring the specific mechanism of how attention operates at the level of information processing, for example by showing that peripheral cues result in a combination of stimulus enhancement and noise reduction (Lu & Dosher 2000). The mechanisms used here would be consistent with both stimulus enhancement and noise reduction

### 3.2 How it works

RAGNAROC simulates the consequences of attentional decisions rippling through the visual hierarchy, creating transient attractor dynamics that allow attention to lock-on (Tan & Wyble 2015; Callahan-Flintoft, Chen & Wyble 2018) to one or more locations. In this context, the term *lock-on* refers to a state in which feedback attentional enhancement from higher-order to lower-order layers of the visual system amplifies feed-forward projections to create a temporary attractor state that anchors attention at a given location for a brief window of time. These lock-on states are similar in some respects to what was originally conceived of as *attentional engagement* (Posner et al. 1987), in that attention is strongly attached to the location of one (or more) stimuli. Importantly, there needs to be something for attention to lock-on to, since the attractor state that drives the lock-on is dependent on feed-forward input. The model also simulates a natural process of disengagement from a given location due to the buildup of inhibition for specific representations at attended locations.

#### Model outputs

The model simulates both the behavioral consequences of reflexive attentional deployment, measured in terms of accuracy and reaction time, while also simulating key ERP correlates of attention, such as the N2pc and the P_D_ components.

#### Stimulus Processing and Differentiation

In order to provide a theory of attention that can be applied to many different experimental contexts, we do not commit to the decoding of pixelwise representations. Instead, simulated neurons in each spatiotopic map represent the presence of attributes at locations with a granularity of 0.5 degrees of visual angle. These representations are segregated into distinct maps that are each specialized for particular kinds of stimuli, as in Itti et al. (1998).

#### Localizing and attending important stimuli

RAGNAROC assumes that attention must determine the precise location of a to-be-attended stimulus from the coarse-grained location information carried by higher levels of the visual hierarchy(DiCarlo & Maunsell 2003), and then deploy attention to the corresponding location.

#### The distinction between Targets and Distractors

RAGNAROC assumes that targets and distractors are distinguished by reflexive attention based on differences in *priority*(defined below), since a decision must be made to commit attention before input from slower, more deliberate stimulus evaluations are completed. In this framework, targets (to the extent that the visual system perceives them as such) are successful at triggering attention because they elicit a stronger priority signal at their location in the attention map. The decision process is, in effect, a race between competing representations, and the top-down attentional set plays a key role in helping task-relevant stimuli to win that race. However, the outcome of this race-based decision process is not pre-ordained and the attention system is prone to deploying attention to highly salient distractors in some cases.

#### Priority value

Each stimulus in the visual field receives a priority value, which is a valuation of its likely importance according to a combination of physical salience, and top-down contributions from attentional set (Figure 3). Physical salience is parameterized to reflect the degree to which a given stimulus stands out from other nearby stimuli in terms of low-level features (e.g. color, orientation, luminance). Priority is also affected by the degree to which a stimulus matches the top-down attentional control settings (Saenz, Buracas & Boynton 2002; Zhang & Luck 2009; Bundesen 1990). These attentional control settings prioritize simple features like color, or more complex attributes such as conceptual categories (e.g. dinner food, animal, etc) by upweighting feedforward activity from some maps and downweighting feedforward processing from other maps. We assume that the ability to select certain stimulus attributes for task-relevant weighting is governed by pre-learned stimulus categories (e.g. contrasting letters vs. digits), but cannot easily be accomplished for arbitrary distinctions (e.g. select letters A, B, C from other letters). This follows from the work of Shiffrin & Schneider (1977) who demonstrated the ability to efficiently attend to previously learned categories, but not to arbitrary subsets of a category. Based on work suggesting that even conceptually defined target sets can be used to select information from RSVP (Potter 1976; Barnard, Scott, Taylor, May & Knightley 2004) or capture attention (Wyble, Folk & Potter 2013), it is assumed that prioritization can happen even at a conceptual level.

**Figure 3.**
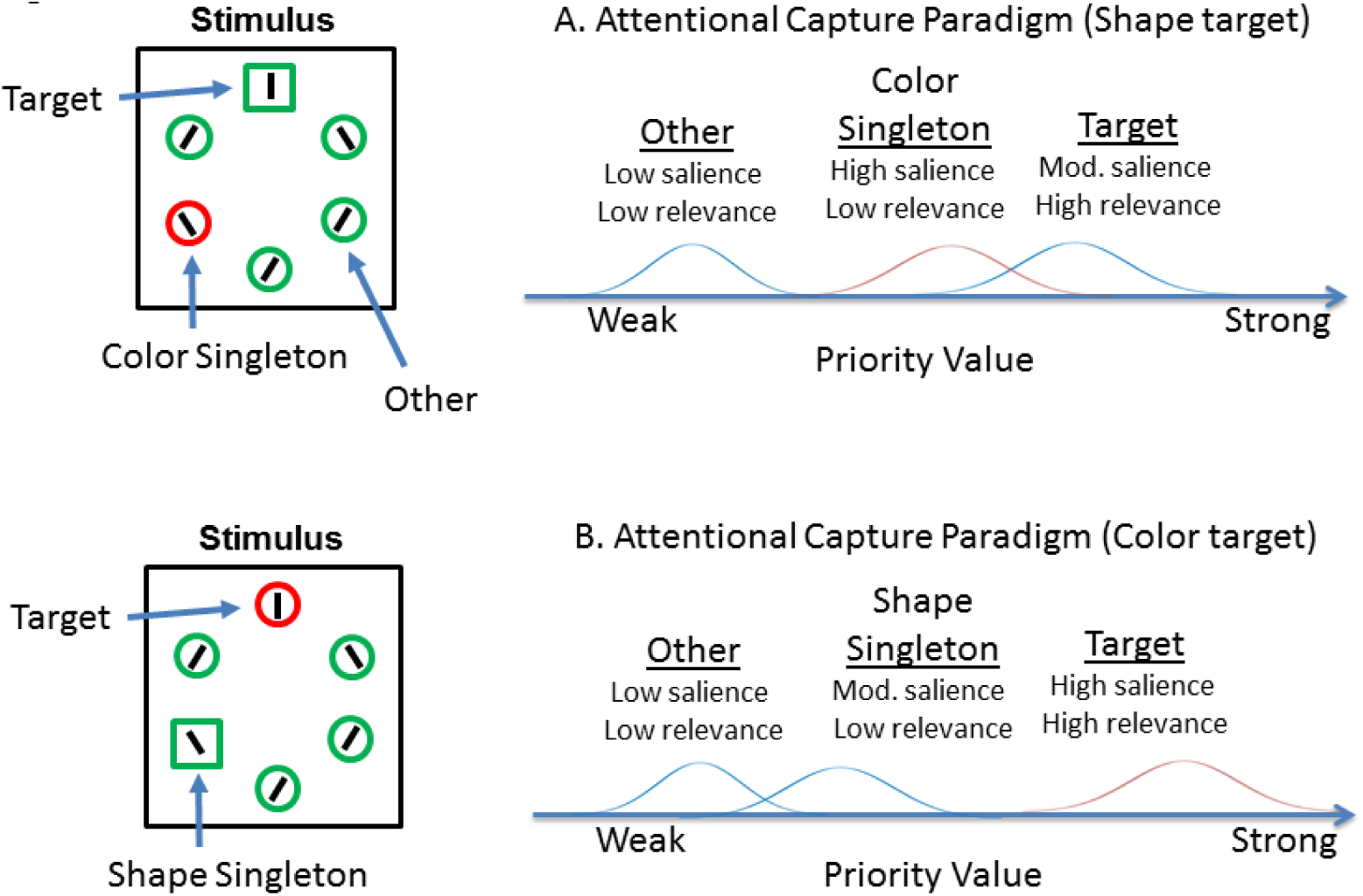
Illustration of how physical salience and top-down relevance can be mapped onto a single priority dimension using canonical attention capture paradigms. A) A highly salient color singleton produces priority values that are competitive with the less-salient target. B) When the target and distractor dimensions are switched, the shape singleton is not competitive with the color-singleton target.

Other potential contributions to priority that will not be explicitly modelled here could involve whether stimulus attributes have been associated with reward (Anderson, Laurent & Yantis 2011), have been recently presented (Awh, Beloposky & Theeuwes 2012), were recently task relevant (Kadel, Feldmann-Wüstefeld & Schubö 2017) or are relatively novel in time (i.e. an oddball). Thus, a strength of the attention-map framework is to allow a broad variety of influences to affect how stimuli are prioritized.

### Attentional Disengagement

Disengagement from a given location occurs rapidly when a location is no longer occupied by a high priority stimulus. Moreover, when a stimulus remains at a given location, an inhibitory process that is stimulus specific leads to a reduction in the strength of attention at that location, such that attention can more easily be attracted by a new stimulus with increasing onset asynchrony. This is caused by inhibitory interneurons (see II neurons in section 3.3.1 below).

#### 3.2 RAGNAROC Architecture

##### 3.2.1 A Hierarchy of Visual maps

RAGNAROC assumes that the visual system is composed of a hierarchy of maps that each represent the visual field and are connected so as to preserve a spatiotopic organization that is rooted in a retinotopic representation at the earliest level (Figure 4). No claims about the number and complexity of this hierarchy are necessary here. Information propagates through the layers via feedforward excitation. The first layer of the model is termed Early Vision (EV) and simulates the earliest cortical regions in the visual hierarchy, which contains neurons with small receptive fields, such as V1. The second tier of layers is collectively termed Late Vision (maps LV1 and LV2) and contains neurons with larger receptive fields, corresponding to anatomical areas in the ventral visual stream that are thought to be specialized for different kinds of stimuli, such as V4 (color), FFA (faces), the EBA (body parts), the PPA (places), as well as distinctions between animate and inanimate stimuli, canonical size (Konkle & Carmazza 2013) and other distinctions that are as yet undiscovered. In our simulations, EV neurons have a simulated receptive field size of .5 degrees, while LV neurons have receptive fields of 3.5 degrees width.

The third layer of the model’s hierarchy is the attention map (AM), which receives convergent input from all of the subordinate LV maps^4^ and has the same diameter of receptive fields as LV neurons. Thus their receptive fields are extremely broad, because their input from the LV is already enlarged. The role of the AM is essentially to implement decision making across the visual field and to enact the consequences of that decision by sending modulatory projections down to the earliest level of the hierarchy. It does this by accumulating spatially imprecise activity from the subordinate LVs and then computing the spatiotopic location of the originating stimulus in the EV by summation. Convergent input from the LV maps initially forms an activation bump, centered at the location of each stimulus in the visual field. The correct localization of this bump follows naturally from the activation dynamics of the model since the AM neuron that resides at the corresponding topographic locus of the centroid of LV activity will receive the largest amount of input from the LV.

When AM neurons are stimulated above threshold, a multiplicative attentional enhancement is applied to feed-forward connections from neurons in the EV at the corresponding spatiotopic location. This modulation increases the strength of the feedforward input from that region of the visual field, which in turn increases the excitatory input to the AM. This dynamic creates a brief attractor state that resonates between different levels of the hierarchy. The activated peak in the attention map is amplifying its own input by enhancing the feedforward projections from the same corresponding spatial location in the EV. This condition we term a *lock-on* state and allows a precise, stable localization of attention despite the relatively coarse-grained spatiotopy in the receptive field of the neurons. The stabilized bump of activation results from satisfying parallel constraints imposed by the recurrent excitation between AM and EV, and the inhibitory gating circuitry within the AM which narrows the spatial focus of the lock-on state.

Thus, the attention map integrates information from the subordinate feature maps to localize one or more targets and then projects an enhancement signal back down to earlier areas at the appropriate location(s). There have been a number of proposals for where such an AM might reside in the brain, including frontal cortex, parietal cortex and portions of the pulvinar nucleus (Shipp 2004). We note that a lateral, parietal location would be broadly consistent with the scalp topography of attention-related ERPs, which are typically larger above parietal cortex than directly over occipital, central or frontal areas (Tan & Wyble 2015) and has a posterior topography (Kiss, Van Velzen & Eimer 2008). It is also possible that this functionality is distributed over several cortical areas, although that would come at the expense of intracortical white matter to mediate the competition.

##### 3.2.2 Attentional gating circuitry

One of the key innovations in this model is the inhibitory control circuitry within the AM (the IG nodes in Figure 4B), which has been developed according to pivotal findings in the literature. It allows attention to rapidly focus at one location, while also selectively inhibiting regions of the visual field that contain other visual stimuli (Gaspelin et al 2015; Cepeda et al. 1998).

To do this, gating neurons (IG in Figure 4b) ensure that inhibition is only delivered to neurons receiving excitatory input from the visual field. Each IG neuron is paired with one principal neuron in the AM. An IG only becomes activated when it receives lateral excitation from another AM neuron (i.e. the curved arrows at the top of Figure 4B) and concurrent excitation from any LV neuron at its own spatial location (i.e. the rising arrow from LV to AM in Figure 4B). As a consequence, inhibition is routed selectively to spatial locations that have information to suppress.

Another key part of the gating circuitry is the disinhibitory component, which increases the stability of lock-on states, such that once a decision to attend to a given location is reached, it is less likely that other stimuli will cause it to disengage. This also allows multiple locations to be attended in parallel, when stimuli are of similar priority. This is accomplished by allowing any strongly active AM neuron to inhibit its paired IG neuron, a form of competitive inhibition that has been determined to have a similar stabilizing function in well-charted nervous systems such as the drosophila larva (Jovanic et al. 2016).

The rationale for using this circuit is two-fold. First, the simpler approaches to attentional suppression that are typically used in such models (either winner take-all, or surround inhibition) are unable to selectively route suppression to a specific location that contains a stimulus while leaving other locations relatively uninhibited. Selective inhibition requires a conjunction of two signals (lateral excitation within the AM and feedforward excitation from the LV) onto the IG neurons in order to generate the targeted suppression.

The disinhibitory component of this circuit (i.e. each AM inhibiting its IG neuron) is crucial for allowing several AM nodes to remain strongly active at the same time once the competition has been resolved. Without this implementation, it was unclear how to obtain an attentional configuration that remained stable when multiple locations were co-active. Moreover, the attention mechanism must be able to both selectively enhance and suppress multiple locations. In the model simulations below, we will illustrate the importance of this circuit by showing how attention to two locations is impaired in the absence of this circuit. Other circuits could potentially achieve the same ends with radically different architectures but this particular configuration achieves a compromise of a robust match to the empirical constraints and simplicity. For other examples of gradient architectures through connected networks see implementations and analyses of dynamic neural fields (Amari 1977; Rougier 2006; Strauss & Heinke 2012) and the MORSEL model (Mozer, & Behrmann 1990)

##### 3.2.3 Free and fixed parameters

The model uses predominantly fixed-parameters according to a set of empirical constraints, which are listed below in section 4. These parameter values are invariant for all of the simulations provided below, except for a subset that vary in order to implement the experimental paradigm of each simulation (e.g. timing and location). There are also two partially-fixed parameters that specify the physical salience and task-relevance (i.e. bottom-up and top-down) weightings of each stimulus type. The term partially-fixed reflects the fact that their relative values are determined by the experimental paradigms. E.g. in simulations of the additional– singleton paradigm (Theeuwes 1991), we allow the specific value of the distractor’s salience to vary, but it has to remain higher than the salience of the target. Finally, there is one additional free parameter that defines the accumulator threshold for a behavioral response for each experiment. This parameter is constrained to have a single value for all conditions of a given experiment and prevents behavioral accuracy values from being at ceiling or floor. The specific values of all parameters are provided for each simulation in the appendix and a full set of code is available here (https://osf.io/rwynp/)

#### 3.3 Mechanisms of the model

##### 3.3.1 Equations

The model uses rate-coded neurons, governed by the activation equations of O’Reilly & Munakata (2001) as shown in the Appendix. In these equations the activation level of each neuron is governed by three currents: excitatory, inhibitory and leak. This level of abstraction is a compromise that captures the properties of synaptic interactions in broad strokes, while allowing rapid exploration of different model architectures. Moreover, the distinction between excitatory and inhibitory currents provides a mapping to current flows underlying EEG components. This set of equations has been used effectively in previous simulations of attentional processes at similar time scales (e.g. Bowman & Wyble 2007).The only exceptions to strict neural plausibility are the use of maximum functions at several points, in order to simplify the neural implementation^5^.

Each connection within the model is enumerated in Figure 5 and their properties are described below. This is a different visualization of the same information shown in Figure 4, but with greater specificity should a reader want to understand the model precisely. Direct projections from the EV to the AM could exist but they are not represented for simplicity.

**Figure 4.**
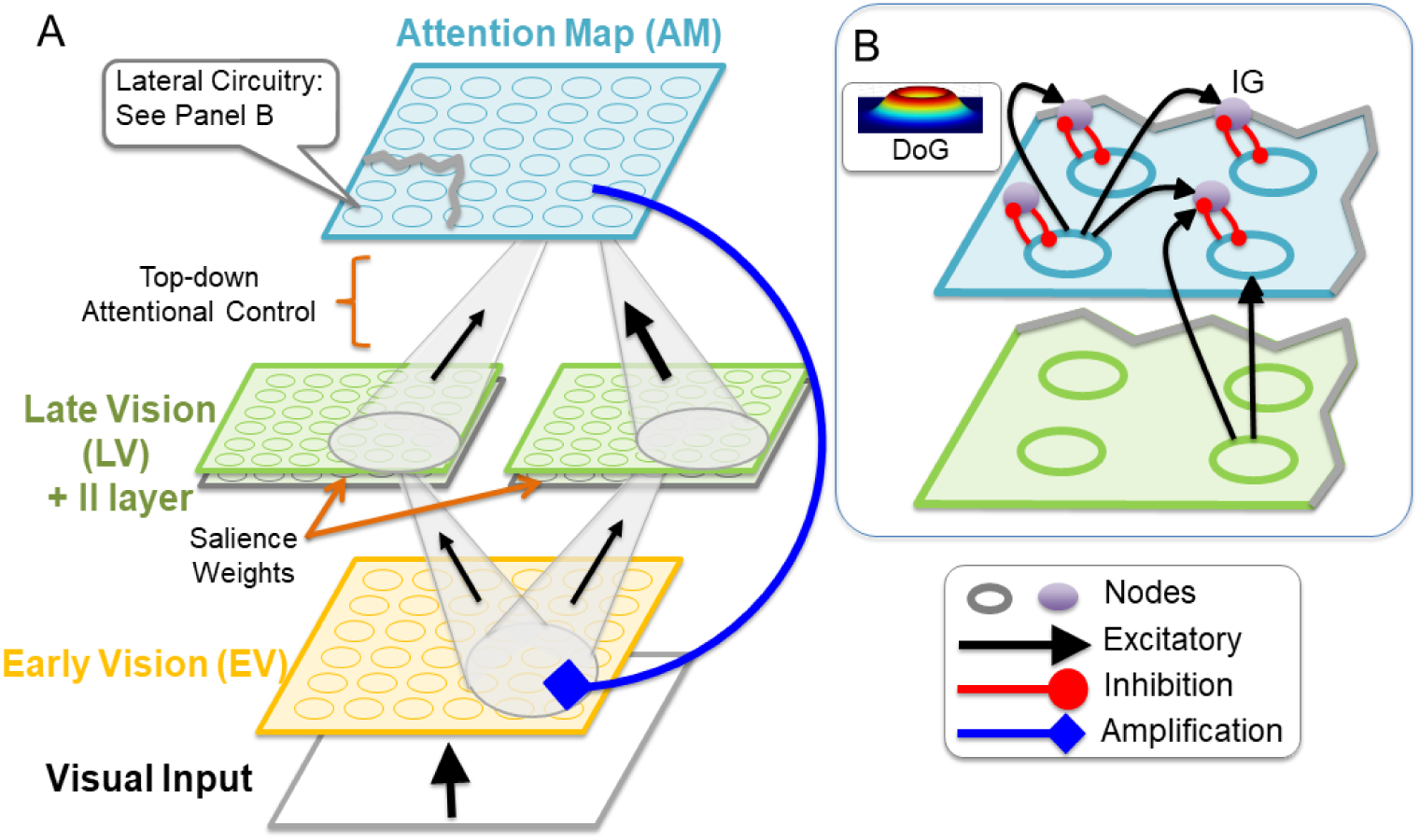
Illustration of the model’s macro architecture (A) and the microcircuitry (B) of the attention map. Figure 5 depicts the same architecture with greater specificity. In the hierarchy of visual areas, the cones reflect the set of neurons at a subordinate layer that excite a given neuron in the superordinate layer. Only two LV maps are shown here, but this architecture generalizes to additional maps. Differences in salience are implemented as stimulus-specific differences in feedforward excitation between EV and LV. Top-down selection is implemented as feature-specific but spatially non-selective modulation of feed-forward weights for an entire LV map. The Attention Map returns location-specific gain enhancement to given locations in the EV. The grey II layer represents feedback inhibition for each LV node. The inset in B shows how neurons are interconnected within the AM. The small grey circles are Inhibitory Gating (IG) neurons, each of which has a competitive inhibitory relationship with a principle neuron of the AM. The principle neurons excite one another with a spatial distribution defined by a Difference of Gaussians (DoG). This connection corresponds to the black curved arrows in panel

For simplicity, all maps have the same dimensionality. Furthermore only two pathways are depicted here though these mechanisms generalize to more complex architectures with more layers and more pathways. All connections between or within layers are assumed to have either an identity projection (i.e. strictly topographic), a Gaussian spread, or a Difference of Gaussians (DoG).

The following numbers indicate connections specified in Figure 5.

**Figure 5.**
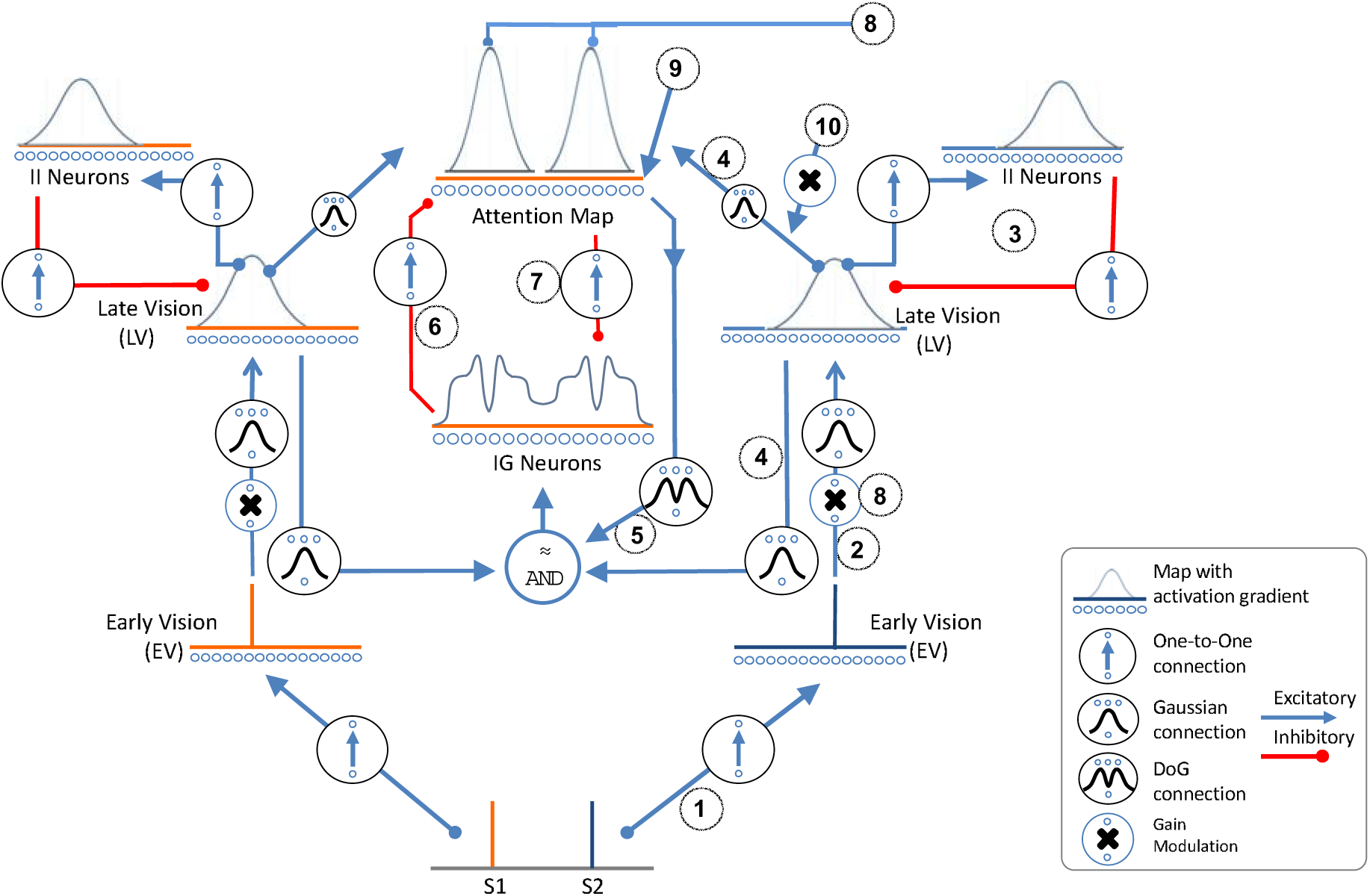
Illustration of the complete architecture for two distinct pathways during perception of two stimuli S1 and S2 with highly distinct dimensions (e.g. form vs color). The left and the right side represent the same connections but for different kinds of stimuli. The bubbles indicate the spatial distribution and character of each connection. The traces above each layer illustrate a typical activation profile for that map in response to a stimulus. Note that attention affects both pathways, regardless of which stimulus triggered it. The numbers correspond to descriptors in the text.

1. Input to EV neurons: The EV represents the earliest stage of cortical visual processing in which neurons have extremely small receptive fields and color/orientation/frequency specific firing preferences. For the sake of simplicity, EV nodes are separated into different areas according to the kinds of stimuli presented, although in the brain these different neurons occupy the same cortical map (i.e. V1). Input to a specific EV node is specified as a step function, since the simulations are of suprathreshold stimuli (i.e. a value changing from 0 to 1 while the stimulus is visible), which causes the corresponding EV node’s membrane potential (MP) to charge up according to equations 1.1,1.2 and 1.3, see appendix.
2. Projection from EV to LV: When an EV node’s membrane potential crosses threshold, it sends a feedforward excitation to an array of nodes in the corresponding LV maps. This projection is spatially weighted according to a Gaussian centered at the location of that EV node. The magnitude of this projection is the salience of the stimulus, and indicates its physical dissimilarity to other stimuli in the visual field according to the specific LV it projects to (e.g. a shape singleton would have a high salience in an LV map that is specific for form). Other accounts have shown how to compute salience for some classes of features such as color, orientation and luminance (Zelinsky 2008; Itti et al. 1998; Bruce & Tsotsos 2006). In RAGNAROC, we abstract over the process of computing salience to accommodate the broad diversity of tasks. Computing LV activation corresponds to equations 1.4-1.6 in the appendix.
3. Inhibitory feedback nodes in the LV: Each LV node has a dedicated inhibitory interneuron (labelled II), that provides feedback inhibition. This inhibitory feedback is crucial for emphasizing the onset of new information by causing the activity of any given LV neuron to drop after approximately 100ms, which is characteristic of single units in the visual system (e.g. Fig 9 of Chelazzi, Duncan, Miller & Desimone 1998). Moreover, these II neurons can cause attention to naturally disengage from a stimulus that remains constant on the retinal field. Computing II activation corresponds to equations 1.8 to 1.10 in the appendix.
4. Projection from LV to AM: When an LV node crosses threshold, it projects feedforward excitation to an array of nodes in the AM according to a Gaussian profile centered at the location of the active LV node. LV nodes also excite IG nodes (see below) with the same Gaussian profile. The projections to both the AM and IG nodes include a parameter that represents the task-relevance (i.e. “top-down”) weighting of each stimulus type, and is fixed for each LV map. (e.g. to represent an attentional set for a specific color, all LV nodes for that color have an increased feedforward strength to the AM). Computing AM activation corresponds to equations 1.11 to 1.14 in the appendix. These connections are depicted as the two ascending black arrows in Figure 4B.
5. Inhibitory Gating Nodes (IG): The IG nodes ensure that inhibition within the attention map only occurs at locations receiving input from an LV (see also Beuth & Hamker 2015). Each IG node is paired with a single AM node that it can inhibit. An IG node is excited by neighboring AM nodes according to a Difference Of Gaussians (DoG) activation profile. IG nodes are also excited by LV nodes. The total excitation of each IG node from these two sources (AM and LV) is capped such that concurrent AM and LV activity is required to raise an IG node above threshold. Thus, IG neurons exhibit the equivalent of a logical AND gate in that they require concurrent activation from two pathways in order to fire. Computing AM activation corresponds to equations 1.16 to 1.21 in the appendix. This connection is depicted as the laterally connecting black arrows in Figure 4B.
6. IG inhibiting the Attention Map: When activated by convergent AM and LV input, an IG node inhibits its corresponding AM node. This is the basis of inhibitory suppression of attention within the model and corresponds to equation 1.12 in the appendix. This connection is depicted as the inhibition from IG to AM in Figure 4B.
7. Attention Map inhibiting IG: This disinhibitory circuit increases the stability of an AM lock-on state, since AM neurons can protect themselves from inhibition generated by neighboring AM nodes. The inhibition from AM->IG has a high threshold of activation, so that an AM node protects itself from inhibition only once a lock-on state has formed at a given location. This inhibition corresponds to equation 1.19 in the appendix. This connection is depicted as the inhibition from AM to IG in Figure 4B.
8. Attentional Enhancement: Each AM node provides a gain modulation of synaptic transmission from EV to LV for all EV nodes at the same location. It is this modulation that creates the lock-on attractor between EV and AM since it allows an AM node to increase the gain on its own input. Note that enhancement of a given location in the EV occurs for the entire Gaussian spread of an EV neuron’s feedforward projection, unselectively across all feedforward pathways. This enhancement corresponds to equations 1.4 and 1.15 in the appendix.
9. AM excitatory Bias: There is a uniform level of bias input to the entire AM, keeping these neurons slightly active in the absence of input. This enhancement corresponds to equation 1.11 in the appendix.
10. Noise input: Intertrial variability is added to the model as modulations of the weights between the LV and AM, which represents fluctuations in attentional control. The variability is constant for a given trial and varies between trials as samples from a Gaussian distribution.

### 3.4 Example Simulations

The following examples illustrate the model’s dynamics in response to several stimulus scenarios.

#### 3.4.1 The simplest case: Single stimulus

When a single stimulus of sufficient priority is presented to the EV, it triggers a *lock-on* of attention at its spatiotopic location, which is a self-excitatory attractor state resonating between EV and the AM through the LV. Figure 6a illustrates the impulse response function in the AM elicited by a single stimulus. Figure 6b illustrates the time course of activation of each of the layers of the model centered at the location of the stimulus.

**Figure 6.**
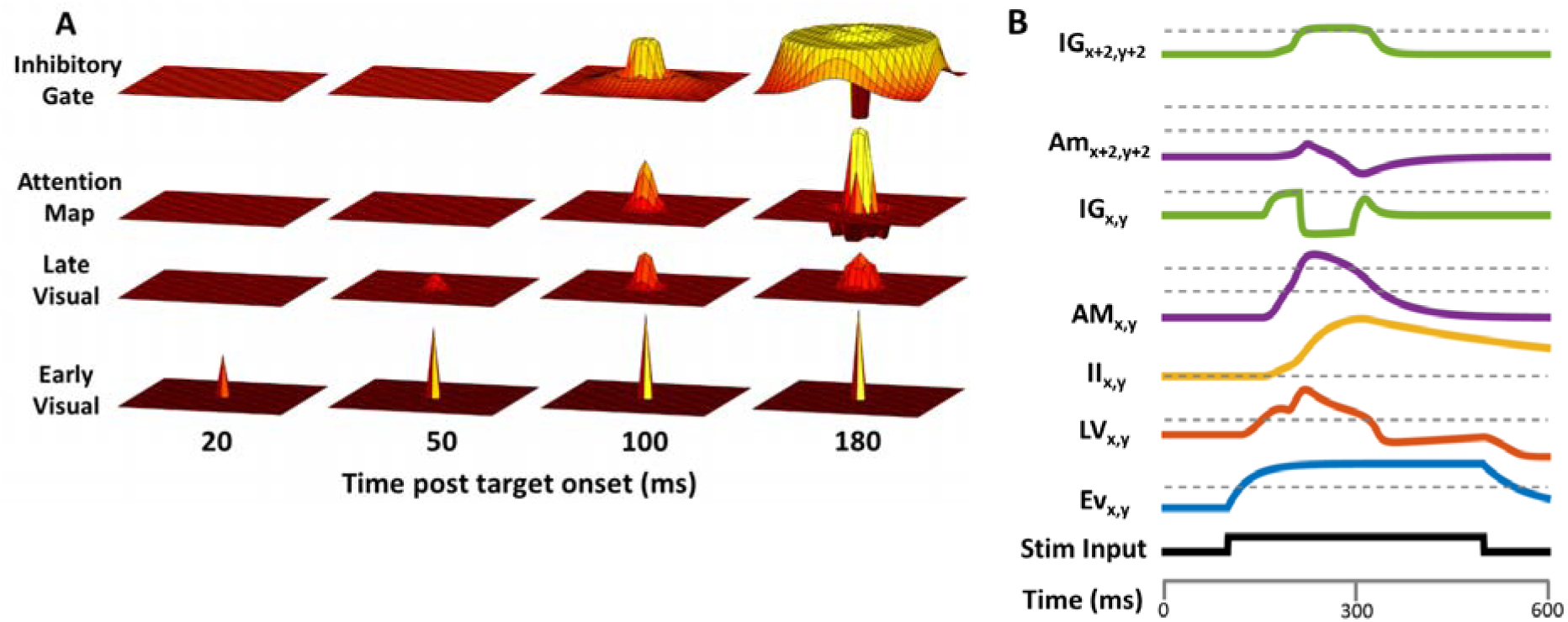
A. Evolution of a lock-on state across time and layers for a single stimulus. An initial feedforward wave of excitation from a single location in the EV triggers activation in the LV, which carries forward to a peak in the AM. Once the central peak of the AM activation crosses threshold, the surrounding IG neurons are activated producing surrounding inhibition in the AM. B. Illustration of the time course of activation for each kind of node within the model in response to a stimulus. Subscripts x,y indicate the location of the stimulus while x+2,y+2 indicate nodes at a neighboring location, 1.4 degree away. Notable inflection points are when the AM_x,y_ node crosses its lower threshold, which triggers enhancement of the LV activation. This drives the AM_x,y_ node more strongly such that it passes its second threshold, allowing it to suppress the IG_x,y_ node. At this point attention is fully locked on to location x,y, since processing is enhanced at that location, and the IG has been inhibited.

#### 3.4.2 Transition to the lock-on state

We demonstrate that the lock-on state has the characteristics of an attractor by illustrating that a broad range of stimulus values evoke a bump in the AM of similar size and duration (Figure 7a). The rapid growth of this neuron’s activation is due to the attentional enhancement of feedforward activity from the EV after the corresponding node in the AM crosses the threshold value.

**Figure 7:**
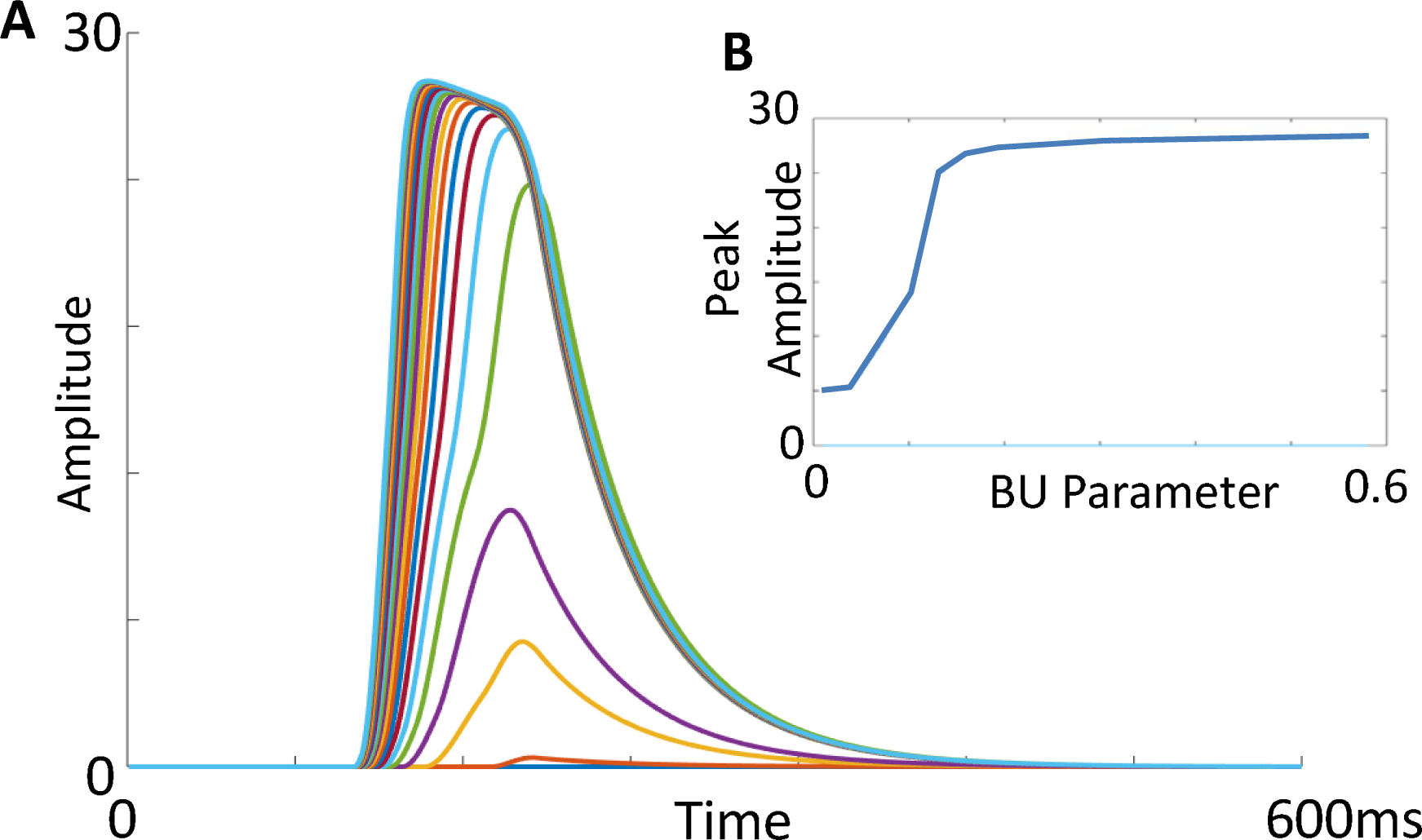
Illustration of the time course of activation of a neuron in the AM centered at the location of a stimulus as the BU parameter is varied from .01 to .6 in increments of .03. The inset plots peak amplitude as a function of activation strength. The takeaway point is that a lock-on state is an attractor, such that many different values of strength map onto the same amplitude of activation.

#### 3.4.3 Two or more simultaneous stimuli

Figure 8 depicts a comparison of AM activity for either one(A), two stimuli(B,C) or six stimuli (D). In the case of two or more stimuli, if one of them is of substantially higher priority, it will inhibit the AM at the locations of the others (B,D). However if both stimuli are of similar priority and nearly simultaneous, they will enter lock-on states simultaneously leading to a fully parallel deployment of attention (C). In such a case, each AM inhibits its paired IG (not shown in this figure), and the lock-on states protect themselves from inhibition by the other.

**Figure 8.**
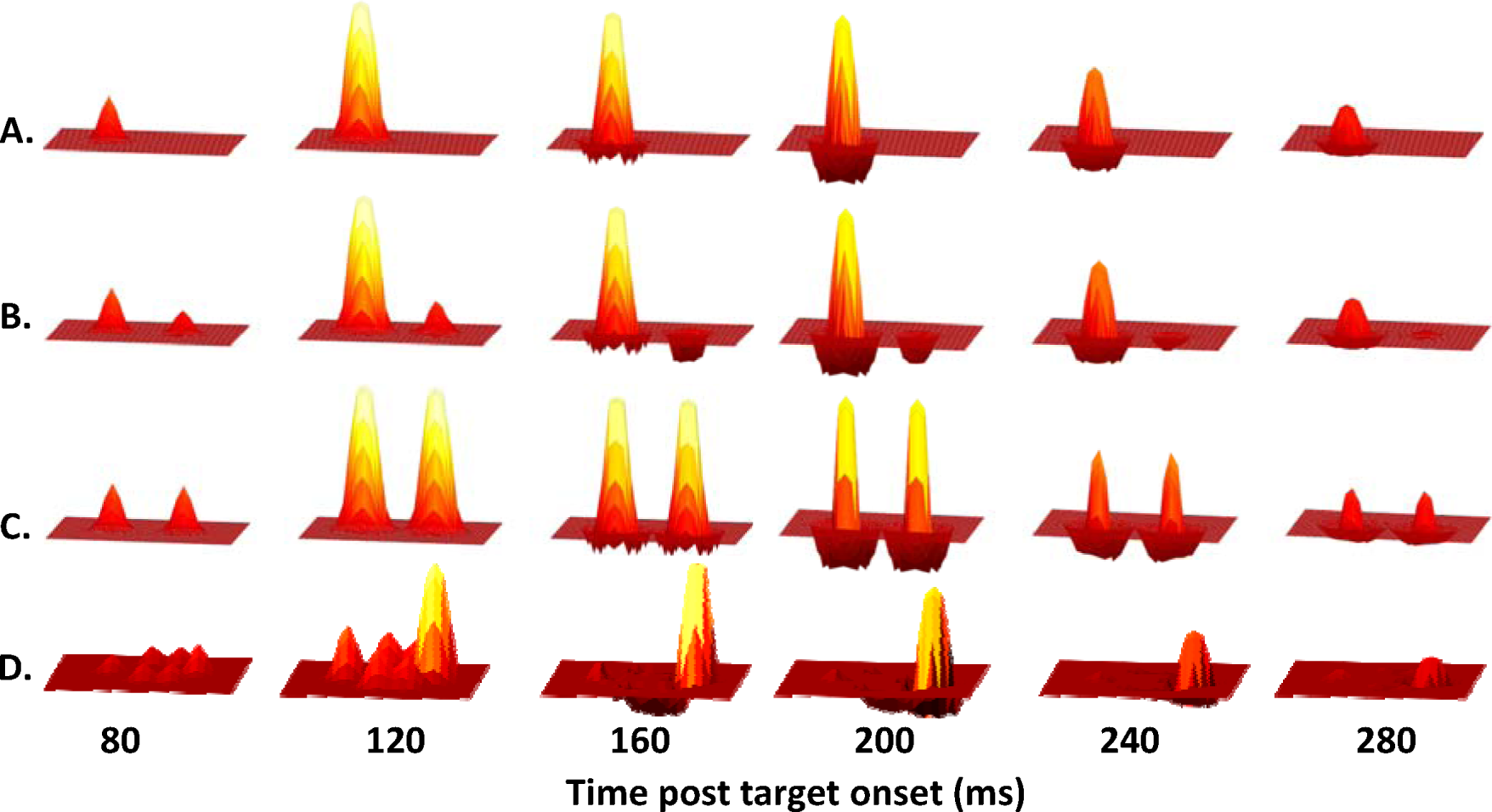
Evolution of activity in the attention map over time for a single stimulus (A), and the same stimulus accompanied by a stimulus of lower priority (B). In the case where two stimuli have equal priority (C), attention is recruited at both locations. (D) illustrates a case in which there is a single stimulus of high priority surrounded by five lower priority stimuli

#### 3.4.4 Two sequential stimuli

When stimuli onset sequentially, the first stimulus will typically be able to activate its lock-on state and suppress activity of the second. In this way, a first target (T1) with a relatively low priority value can suppress attention to a second target (T2) since the temporal advantage of T1 allows it to establish a lock-on state before the T2 has a chance to establish one. To demonstrate the temporal dynamics of how lock-on states interact for two stimuli with varying SOA and priority, Figure 9 illustrates graphically the relative robustness of such states. In the bottom right, the U shaped function illustrates that a T2 can more easily establish a lock-on state if it is presented either simultaneously with, or at least 100ms after a T1. At SOAs near 50ms, the T2 lock-on state is not established unless the T2’s priority is sufficient to countermand the inhibition caused by T1. The bottom left quadrant shows that T1 is not greatly affected by T2’s lock-on state. The top two panels will be discussed in the next section.

### 3.5 Competitive Inhibition helps to stabilize attentional focus

Competitive inhibition improves the functionality of attention by increasing the stability of one or more lock-on states, allowing attention to be simultaneously deployed more easily, in accord with empirical findings such as Bay & Wyble (2014) and Goodbourn & Holcombe (2015). To illustrate the effect of competitive inhibition on the attentional state, Figure 9 compares the intact model (bottom two panels) to one in which the AM->IG inhibition has been removed (top two panels). This change preserves the center-surround inhibition, and the selective inhibition but does not allow AM nodes to competitively block their own inhibition.

With the inhibition intact (bottom two panels), it is much easier for simultaneously (or nearly so) stimuli to evoke robust lock-on states because each protects itself from interference by the other. However within about 50ms, the window of attentional simultaneity has expired because the T1 lock-on starts to inhibit nearby activity in the AM. This makes it difficult for T2 to establish its own lock-on state if it onsets between 50 and 100ms after the T1. The time course of this transition from simultaneous to sequential attention is in agreement with behavioral data showing the onset of attentional inhibition following a T1 onset (Mounts 2000, Experiment 2).

The advantage of protecting the lock-on state of the T1 at the expense of the T2 is to reduce the volatility of attentional decisions in the case of dynamic or rapidly changing stimuli.

In contrast, when the competitive inhibition is removed (top two panels), the AM activations evoked by two stimuli always compete against one another, such that only T1 or T2 can be fully attended. This also allows a delayed onset T2 to interfere strongly with T1, in contrast to typical findings of attention paradigms in which the T1 has strong priority over T2 (Duncan, Ward & Shapiro 1994).

### 3.6 Mapping model activity onto measurable data

In order to compare the model against empirical benchmarks, it is necessary to map model activity to behavioral measures of accuracy and reaction time, as well as EEG correlates of attention such as the N2pc and P_D_. Figure 10 illustrates which activity states in the RAGNAROC model are used for generating data.

**Figure 9.**
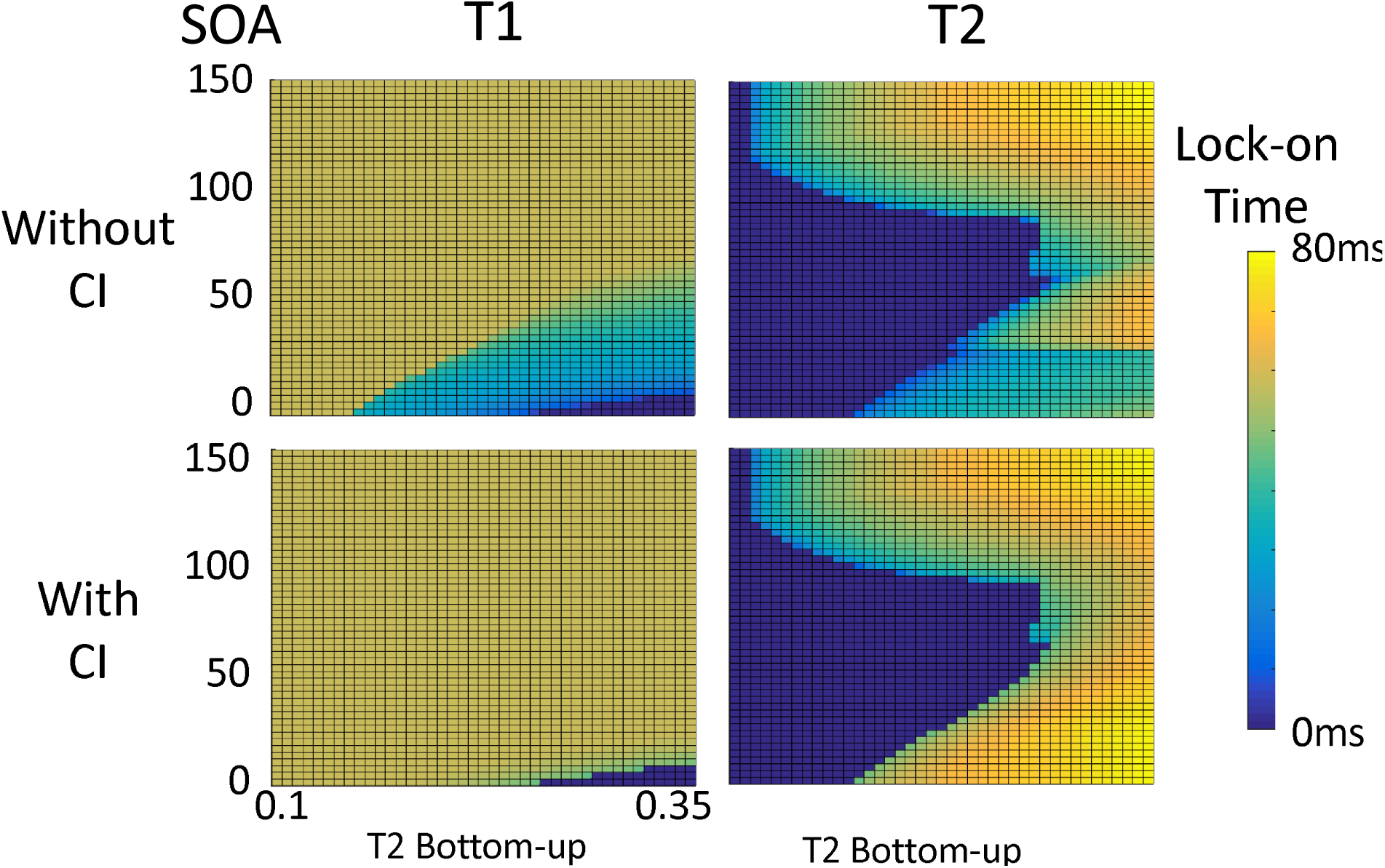
Illustration of the robustness of lock on states to each of two stimuli where the second stimulus varies in the strength of bottom-up activation parameter of the T2 (horizontal axis) and temporal latency (vertical axis). Each point represents the total duration for which the AM neuron at the location of the stimulus (T1 or T2) is above threshold. The key takeaway is that without competitive inhibition (CI), T1 and T2 compete destructively at short SOAs such that neither elicits a robust lock-on state. With inhibition intact, both stimuli can achieve a lock-on state if presented with nearly identical onsets. However at longer SOAs T1 suppresses T2 enforcing a serialized deployment of attention. In this simulation, T1 and T2 were presented 4 degrees apart and have a duration of 120ms. Note that the T1 bottom-up weighting is fixed for all simulations and only the T2 weighting is varied. The distinction between T1 and T2 becomes notional when they are simultaneous.

**Figure 10.**
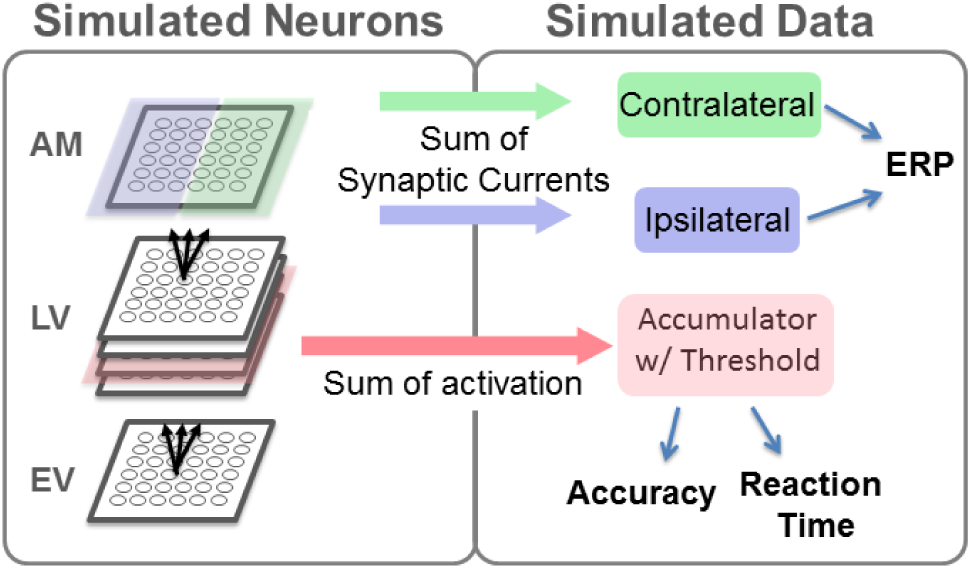
Activity within the RAGNAROC model is used to construct simulations of data in the form of behavioral accuracy, behavioral reaction time and EEG components. Behavioral data are extracted from the late vision area, which is assumed to drive the formation of memory representations and response decisions through mechanisms that are outside of the scope of the model. Simultaneously, synaptic currents within the attention map are measured to generate simulated ERPs such as the N2pc and the PD.

#### 3.6.1 Model configuration

For each experiment, physical salience values and task-relevance weightings are configured for different kinds of stimuli in the task. To provide variability, task-relevance weightings are varied at random over repeated simulations, while the physical salience values remain fixed. Task-relevance weights are initially varied according to a uniform distribution of possible values for each kind of stimulus in the task. For example in a salient-singleton attentional capture paradigm (Theeuwes 1992), the target stimulus has a physical salience value of .15 and a range of task-relevance weightings from .17 to .37 in 12 steps of .018. The singleton distractor has a physical salience of .3 and a range of relevance weightings from .07 to .27 in 12 steps of .018. The model is run for all possible combinations of these weightings, (e.g. 144 total simulations in this example). The simulations are then bootstrapped to form the simulated data set of an entire experiment. The bootstrap involves resampling the model output 1,000 times according to a normal distribution projected onto the 12 TD weight parameter values for each feature dimension, with endpoints of +4/-4 standard deviations under the assumption that there is trial to trial variability in the attentional set of the observers that is normally distributed. This bootstrapping determines both the simulated behavior (accuracy and RT, as appropriate) and the EEG traces for a given experiment. This is the only source of variability in the model when simulating EEG. For simulations of behavior, an additional source of noise is added during the bootstrapping, as described below.

#### 3.6.2 Simulating Behavior RAGNAROC simulates the successful detection or response to a target with a thresholded accumulator

The accumulator sums the time course of activation of all LV neurons that are activated over baseline (.5) for the target. For every trial, the area under the curve (AUC) is calculated for the entire time course of activation. A trial is considered accurate if this AUC exceeds a threshold value that is calibrated for each task. This threshold is a free parameter fit for a given task to achieve a particular accuracy value in one baseline condition chosen for each task. In order to introduce more variability and make simulations less sensitive to the particular threshold value, each simulated trial is jittered with random noise. This is done by adding 15% of the baseline condition’s average AUC times a random scalar (ranging from 0 to 1 from a uniform distribution) for each trial. Reaction times are calculated as the time step at which the accumulator crossed threshold.

#### 3.6.3 Simulating EEG

To translate from simulated neurons to EEG correlates is a hard problem that, in its most exact form, would require compartmental-level modeling of cortical neurons including all of the synapses in each layer, a fairly complete understanding of the neuroanatomy for each individual subject, and a model of the electrical properties of the tissue layers above the cortex (dura, fluid, skull, muscle, skin).

However, it is possible to make effective progress with a much simpler model, given some starting assumptions to simplify the forward model for generating scalp potentials. Here, these assumptions are (1) that the attention map exists over a region of cortex situated in posterior-lateral parietal areas (2) that EEG potentials are largely driven by excitatory synaptic input on pyramidal neurons oriented perpendicular to the scalp (Nunez & Cutillo 1995) (3) that an increase in this synaptic current within the attention map produces (on average) a negative voltage at the scalp and (4) that there is an additional weighting parameter that determines the relative contribution of excitatory and inhibitory synaptic currents for all simulations. The advantage of such a simple model is that it provides fewer opportunities to overfit the observed EEG.

Given these assumptions, RAGNAROC simulates lateralized EEG components associated with attention by summing synaptic currents across each half of the AM, and taking the difference of those sums relative to the side of the visual field that a particular stimulus was presented on.

This is analogous to the measure of potentials such as the N2pc and P_D_, which are calculated as the difference in voltage between electrodes contralateral and ipsilateral to the side of the display containing a target (or a distractor in some cases). Simulations with multiple trials in a block compute the EEG as an average across the trials, and a Gaussian smoothing operation is then applied with a 50ms window.

The synaptic currents are computed separately for each neuron as its excitatory current, minus its inhibitory current, with a floor of zero. The intuition behind this implementation is that excitatory currents are the primary drivers of the large dipoles that are observable at the scalp, and inhibitory inputs often shunt those excitatory currents by creating high conductance areas of the cell membrane closer to the soma (Koch, Douglas & Wehmeier 1990).

The AM receives a uniform input to elevate all of the neurons above their resting potential. This provides a baseline level of excitatory current that is uniformly distributed across the attention map and therefore drops out during the subtraction of ipsilateral from contralateral. Activation or inhibition of nodes within the AM causes deviation away from this baseline level of current.

When this current is summed across the halves of the attention map, laterally asymmetric differences in activation produce changes that are comparable to the N2pc and P_D_ components ^6^. Note that using the sum of currents across a hemifield to simulate voltage means that increases in current for some neurons might be effectively invisible to the simulated EEG signal if there are also corresponding decreases in current for other neurons in the same half of the attention map (Figure 11). Furthermore, any negativity in the simulated voltage difference between the contralateral and ipsilateral sides of the map could be caused either by an increase in activity in the contralateral side, or a decrease in activity on the ipsilateral side. It is important to remember that there are only two possible polarities of a component, positive or negative, but there are (many!) more than two neural processes that could result in a scalp potential at a given latency. Therefore one cannot uniquely ascribe a given functional property to an ERP on the basis of a given polarity/latency.

**Figure 11.**
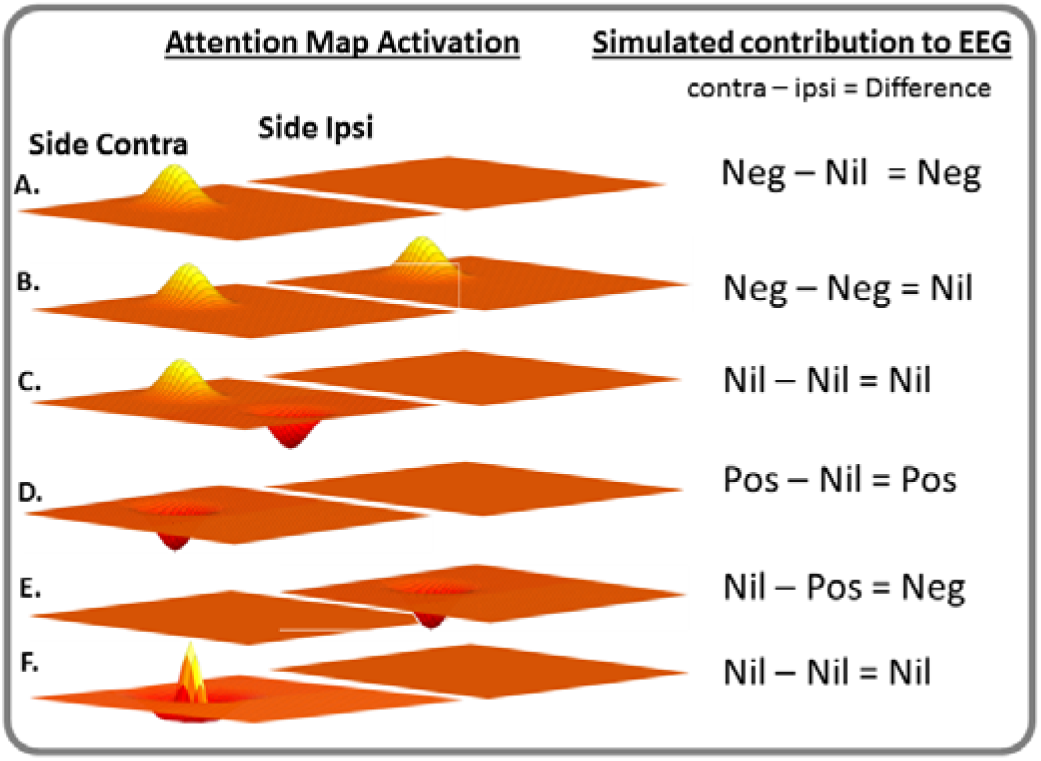
Illustration of how different patterns of activity on ipsi and contralateral sides of simulated cortex can summate to produce either positive, negative or nil voltage differentials. Note that in F, the currents generated by the activity in the peak is effectively cancelled out by the surrounding inhibitory surround producing an effective Nil in the contralateral side. Note that “Nil” in this context doesn’t necessarily mean exactly 0, but sufficiently small that it is not detectable at a given level of experimental ower.

This ambiguity in the interpretation of simulated EEGs is not a shortcoming of the model, but rather reveals a complication inherent in the interpretation of all ERPs. This complication underscores the importance of understanding EEG signals at the level of their neural sources and the role of computational models in understanding those sources.

#### 3.6.4 Simulation of the N2pc

In RAGNAROC, any lateralized stimulus that has the highest priority produces a simulated ERP that resembles an N2pc (Figure 12a). The onset and peak of the N2pc is caused by the initial activation bump in the AM. When the lock-on state is established, the AM activates its neighboring IG neurons, which adds an inhibitory region in the immediate surround. When the central peak is surrounded by inhibition, the sum total of synaptic currents on the contralateral side of the AM nearly cancel out and sometimes even reverse briefly producing a positivity.

**Figure 12.**
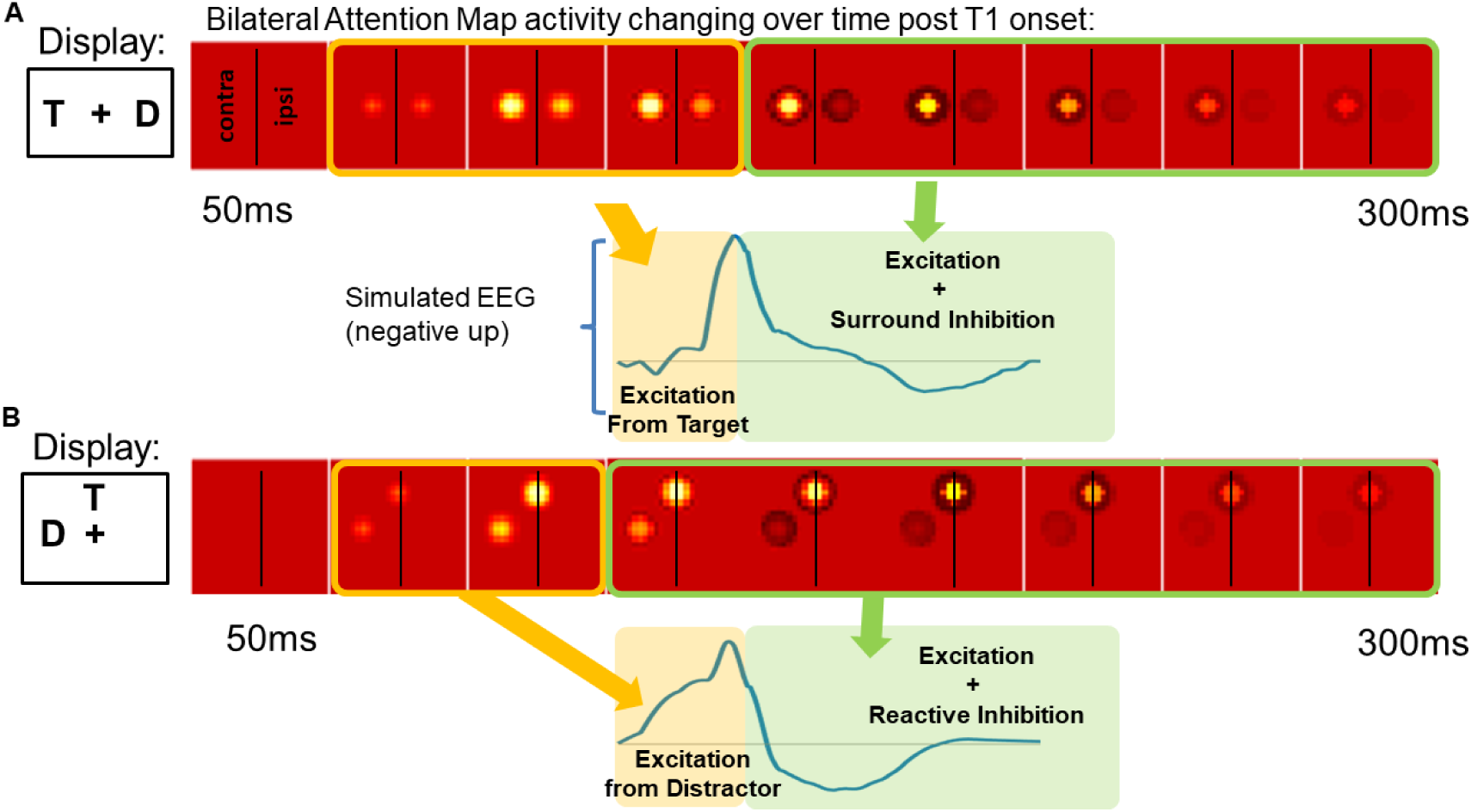
Illustration of how activation levels within the attention map produce simulations of N2pc and Pd components for commonly used experimental paradigms with targets (T) and distractors (D). Note that this is not intended to predict that distractors always elicit an N2pc.

Thus, while RAGNAROC is in general agreement with the theory that the N2pc reflects processes associated with spatial attention, it suggests a more specific temporal relationship, which is that the N2pc reflects, in large part, the processes of localizing a target prior to attentional deployment, as in the CGF model (Tan & Wyble 2015; Callahan-Flintoft, Chen, & Wyble 2018).

Thus, the model explains that the end of the N2pc does not indicate the termination of attention, but rather the onset of surround suppression. This simulation provides a straightforward explanation for the duration of the N2pc, which is typically brief and followed by a positive rebound (Brisson & Jolicouer 2006)

#### 3.6.5 Simulation of the P_D_

The P_D_ is an EEG component thought to reflect inhibition of distracting information in the visual field. In RAGNAROC, a P_D_ can emerge whenever there is sufficient inhibition of activity in the attention map and this occurs in at least two ways. First, whenever two stimuli compete for attentional control and one of them loses, the AM is suppressed at the location of the loser (Figure 12b). This suppression reduces synaptic currents in the hemifield containing that stimulus and results in a net positivity in contralateral scalp electrodes. However, a P_D_ also occurs when the surround suppression encircling an attended stimulus is large enough that it causes a net reduction in current for that half of the visual field. This imbalance would be reflected as a P_D_ trailing an N2pc, and could occur even in the absence of a suppressed distractor (Figure 12a, see also Töllner, Zehetleitner, Gramann, & Müller 2011).

These are the essential aspects of simulating behavioral effects as well as lateralized EEG components in the early time range following the onset of a stimulus array. In the next section we illustrate how specific empirical effects emerge in specific experimental contexts through these mechanisms.

## 4. Simulations of Empirical Constraints

### 4.1. Constraints in model development

As in previous papers (Tan & Wyble 2015; Swan & Wyble 2014), the model is parameterized according to a set of extant findings in the literature. Once the model is able to simultaneously accommodate those findings with one set of fixed parameters, it can be used to generate insights about the underlying system and testable predictions for future work. The philosophy of this approach is to allow a large number of empirical constraints to inform the model’s design, with as little parametric flexibility as possible. Here we list a series of behavioral and electrophysiological findings that we consider to be crucial for defining the functionality of reflexive attention. Each of these findings is simulated with the same set of parameters, except for the configural parameters described in 3.2.3. The supplemental describes the exact set of parameters for each simulation. Given the large diversity of experimental paradigms that provide the constraints, the fits are evaluated for their qualitative similarity to the data.

### 4.2 Behavioral constraints

1. Covert spatial attention is triggered rapidly in response to a target or highly salient stimulus. This effect is measurable as an enhancement of accuracy and reduced reaction time for stimuli presented just after a cue, at that same location. The time course of this enhancement peaks at about 100ms SOA (Nakayama & Mackeben 1989). Note that this transient form of attention is short lived even when the cue stays on the screen. It is difficult to precisely estimate the duration of this effect because it is followed by slower, more volitional forms of attention that sustains the attentional effect to differing degrees in differing paradigms. However, there have been consistent findings of enhanced perception at brief cue-target (Yeshurun & Carrasco 1999; Müller & Rabbitt 1989; Cheal & Lyon 1991) or target-target intervals that attenuate at longer cue-target intervals. Targets elicit such attention as well (Wyble, Potter, Bowman 2009). RAGNAROC simulates the transient attention effect of Nakayama and Mackeben (1989) as a brief window of elevated accuracy in reporting a target when it follows another stimulus at the same location because the second stimulus benefits from the lingering lock-on state created by the first stimulus. (Figure 13a, the two traces in the data plot indicate different subjects)
2. This rapid deployment of attention is reflexive, which means that it is vulnerable to capture by a non-target stimulus that is either highly salient (Theeuwes 1992) or contains a target-defining attribute (Remington, Folk & Johnston 1992). This reflexive form of attention occurs even to locations that are known to always be task irrelevant (Lamy, Leber & Egeth 2004; Krose & Julesz 1989). Also, highly-salient distractors will trigger this form of attention regardless of instruction, or lengthy practice sessions (Theeuwes 1992; but see Kim & Cave 1999 for a counter example). RAGNAROC simulates the attentional capture effect of Theeuwes (1992) as a longer reaction time for a target in the presence of a distractor (Figure 13b). See the discussion section for an in depth discussion of precisely what causes the slower RTs in a capture paradigm.
3. Reflexive attention can be biased towards stimuli containing certain features or attributes, provided that there exist well-learned, cognitively accessible distinctions between target-defining features and other stimuli (e.g. letters can be selected among digits but an arbitrary subset of letters cannot be efficiently selected from other letters without substantial training, Schneider & Shiffrin 1977). This target-defining attentional set is implemented across the entire visual field such that, for example, establishing a control setting for red at one location prioritizes red at all locations (Zhang & Luck 2009). RAGNAROC simulates attentional set as capture costs that are mediated by task-set from Folk, Remington & Johnston (1992). See Figure 13c.
4. Reflexive attention can be deployed to two or more locations at the same time when stimuli are presented in parallel, but behaves more like a spotlight when targets are presented sequentially (Bichot et al. 1999; Dubois Hamker & VanRullen 2009; Bay & Wyble 2014). RAGNAROC simulates divided attention as an attentional benefit that is similar in size regardless of whether one or two locations are cued (Bay & Wyble 2014). See Figure 13d.
5. Presenting a cue or target at one location causes subsequent targets presented at spatially proximal locations to be harder to perceive. This suppression is diminished with increasing spatial distance (Mounts 2000; Dubois et al. 2009). RAGNAROC simulates attentional suppression surrounding an attended region using two sequential targets as in Mounts (2000). See Figure 13e.
6. In the presence of a target, inhibition is localized at the spatiotopic position of non-target stimuli, in comparison to empty locations in the visual field. Thus, probe stimuli will be harder to perceive when they occur in the locations of singleton distractors in comparison with blank areas (Cepeda et al. 1998) or non-singleton distractors (Gaspelin, et al. 2015; Figure 2c). Moreover, this effect is dependent on the attentional set of the subject. It is present only when targets are defined by specific features, rather than by being a form singleton. RAGNAROC simulates increased suppression of attention at locations containing salient distractors when the top-down weightings from LV->AM for the target are increased (Figure 14 bottom two panels). When these weightings are weaker, the reverse pattern is obtained such that salient distractors evoke attentional enhancement rather than suppression (Figure 14, top two panels).

### 4.3 EEG Constraints

Select data concerning the N2pc (also referred to as the PCN by Tollner, Muller & Zehetleitner 2012) and P_D_ components will be taken as constraints on the model as well.

**Figure 13.**
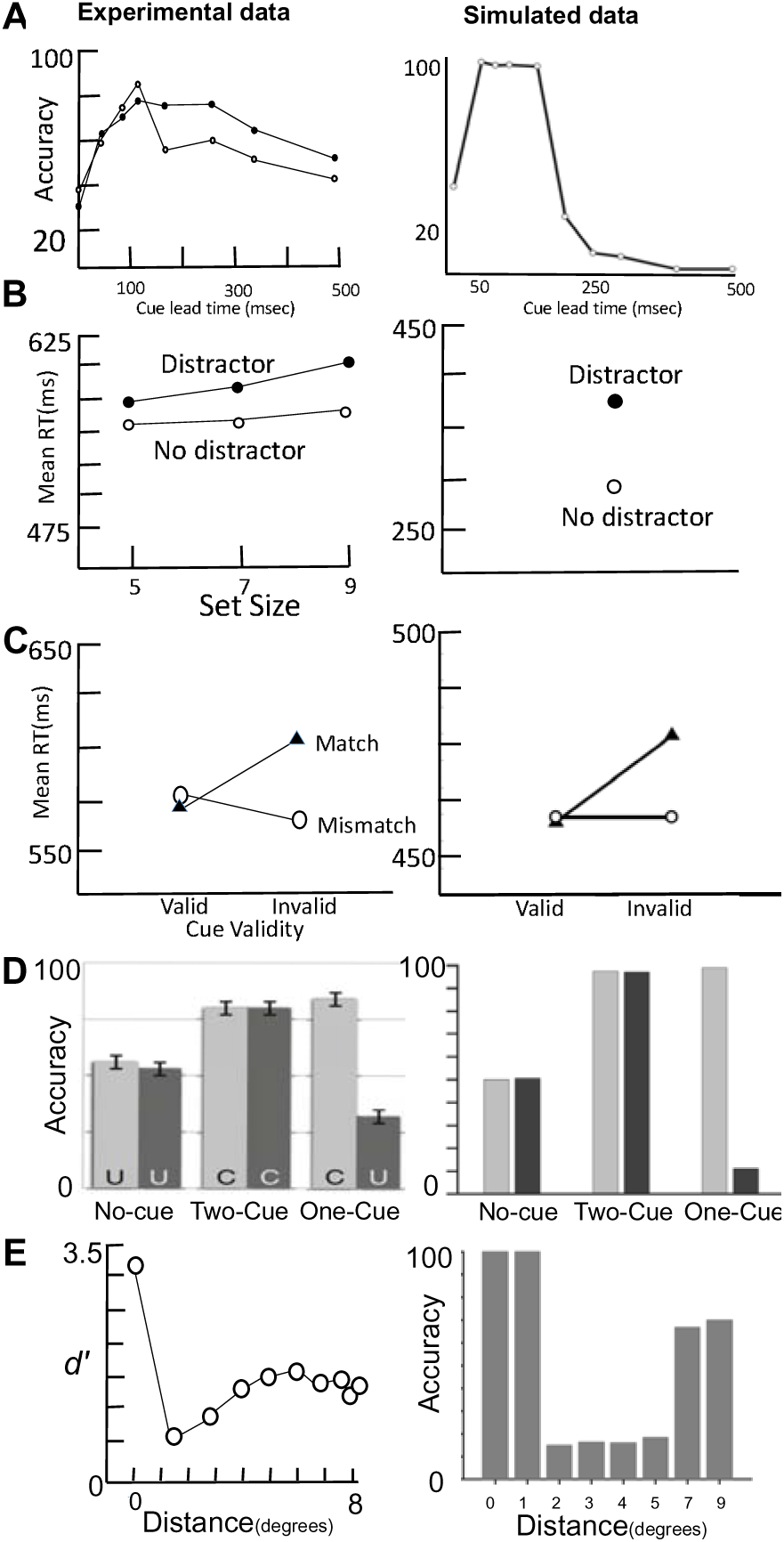
Behavioral constraints and simulations. (a) Accuracy of reporting a target indicates the transient nature of reflexive attention (N = 2, Nakayama & Mackeben 1987). (b) Reaction time to report a shape singleton target is increased in the presence of a salient color distractor (Theeuwes 1992; Experiment 3). (c) A singleton affects the reaction time to report a target only if it matches the type of target (Folk Remington & Johnston 1992; Experiment 3). (d) The benefit of a valid cue, relative to a no-cue condition, is not diminished when two cues are used, suggesting simultaneous deployment of attention to two locations with minimal cost (Bay & Wyble 2014). (e) Accuracy of reporting a second target is affected by proximity to a preceding target with a spatial the empirical data is reported as d’, but model accuracy is reported as accuracy since it lacks a mechanism for guessing.

**Figure 14.**
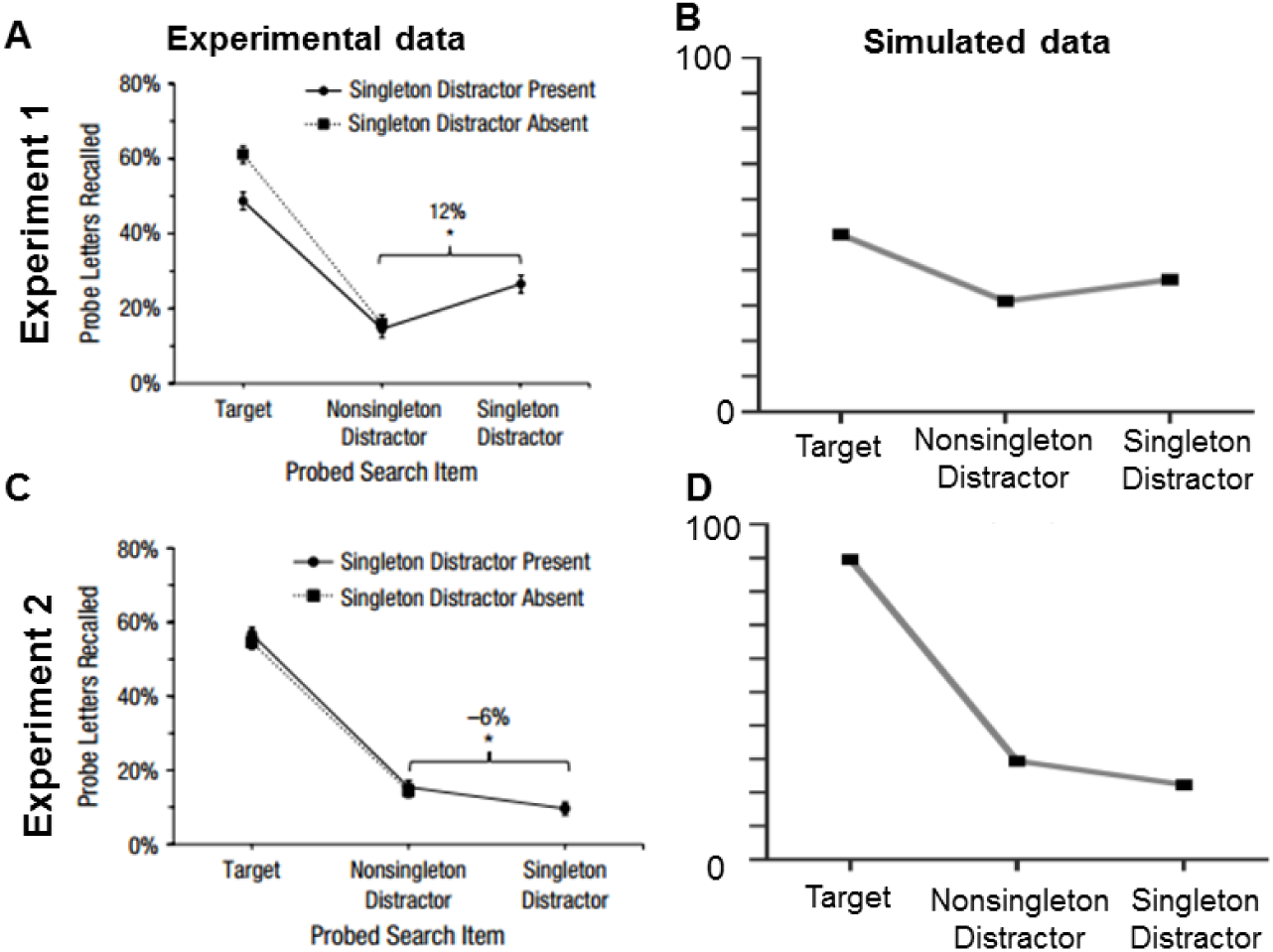
Experimental data from Gaspelin, Leonard Luck (2015) alongside simulations. In panel A the target was a shape singleton And report of the probe at the singleton color distractor was elevated, compared to the nonsingleton distractor. The model simulates this effect (B) as the result of weaker top-down attention which allows the salient distractor to trigger attention. In panel C, the participant knows exactly what shape will contain the target. This is simulated (D) by adopting stronger top-down settings, which allows the target to inhibit the distractor on nearly every trial.

#### 4.3.3 Specific EEG Constraints

1. Presenting a target in either hemifield produces a brief negativity in EEG recorded on the contralateral, posterior side of the scalp called the N2pc (Luck & Hillyard 1994; Eimer 1996) or PCN (Töllner, Zehetleitner, Gramann & Müller 2011). This negativity typically peaks at about 250ms after target onset and is observed even in the absence of distractors on the same side of the display (Tan & Wyble 2015). RAGNAROC simulates this effect as a contralateral negative voltage for a target on one side of the visual field (Eimer 1996). See Figure 15a.
2. Multiple targets in the same location in immediate succession will produce a standard N2pc only to the first target in the sequence, even for trials in which both targets were reported. (Tan & Wyble 2015; Callahan-Flintoft et al. & Wyble 2017; Callahan-Flintoft, Chen, & Wyble 2018). When the two targets are presented in different locations of the visual field, there will be an N2pc to each of them in turn (Tan & Wyble 2015). This single-N2pc effect is only present when the two targets are presented within roughly 150ms and at exactly the same location. When the duration between targets is extended, a second N2pc is observed for the second target, even when it is in the same location as the first and also regardless of whether subjects know that the second target will appear in the same location as the first (Callahan-Flintoft, Chen, & Wyble 2018; See Figure 15c).
3. Multiple N2pcs can be evoked in rapid succession (e.g. at 10-100ms intervals), with no delay when targets are presented at different locations. When presenting a lateral target at intervals of 10, 20, 50 and 100ms relative to a preceding target, an N2pc is evoked with a target-relative latency that is very similar (i.e. within 10ms) to that evoked by the first target. This finding indicates that deploying attention to one target does not affect the time course of attentional engagement to a second target within this short time frame (Grubert, Eimer 2014, Experiment 1). At longer separations, an attentional blink may be observed but the blink is not within the scope of the mechanisms of RAGNAROC See Figure 15d.
4. The N2pc/PCN is often followed by a positive contralateral potential called the P_D_ (Hickey et al. 2009; McDonald, Green, Jannati & DiLollo 2012) or Ptc (Hilimire, Mounts, Parks & Corballis 2010). This positivity has been particularly associated with the occurrence of a highly salient lateral distractor, although this positivity can occur without such a distractor (Töllner et al. 2011; Hilimire, Hickey & Corballis 2011). RAGNAROC simulates this effect as a positive voltage after the N2pc for a lateralized target (Töllner, et al. 2011). See Figure 15e.
5. A laterally presented salient distractor can produce an N2pc, and this N2pc will be larger if the distractor is presented without an accompanying target (Kiss, Grubert, Petersen, & Eimer 2012; Hilimire, Hickey & Corballis 2012; McDonald, et al. 2012). RAGNAROC simulates this effect as a negative contralateral voltage after a lateralized distractor (McDonald et al. 2012). See Figure 15f.
6. Specificit y of the attentional set affects the degree to which targets and distractors produce N2pcs. When the task set does not predict a specific stimulus (e.g. when the task is to find the shape singleton), the distractor can elicit an N2pc (Hickey, et al. 2006; Burra & Kerzel 2013) because the top down weighting is less efficient, which allows a salient distractor to have higher relative priority. Furthermore, a target presented on the midline will reduce the distractor induced N2pc by competing with it for attention (Hilimire, et al. 2011; Hilimire & Corballis 2014; Figure 3c). Similarly, in the same condition an N2pc induced by a lateral target is reduced by a centrally presented distractor (Hilimire & Corballis 2014). When the task set is a *predictable singleton*, then distractors produce a much weaker N2pc and the target induced N2pc is barely affected by the presence of a salient distractor. RAGNAROC simulates this effect as a specific ordering of N2pc amplitudes for different stimulus configurations across two different specificity manipulations (Hilimire & Corballis 2014). A similar result is obtained when the task is manipulated such that the distractor is of higher or lower salience than the target. For example, when the task is to detect a form singleton, a highly salient color singleton will reduce the target’s N2pc, but this is not true when the task is to detect a color singleton, and the distractor is a form singleton (Schubö, 2009). See Figure 15g.

#### 5.0 Discussion: what have we learned?

The RAGNAROC model describes a set of neural mechanisms that explicates how attention reflexively responds to new visual input, and makes rapid decisions about which locations in the visual field to enhance and which to suppress. The decisions are mediated by attractor states and competitive inhibition that help to ensure that the decisions are stable and accurately targeted at the correct location. It is argued that this reflexive attentional system plays a key role in many experimental paradigms, and constitutes the first form of decisive filtering of visual information after it enters the brain.

**Figure 15.**
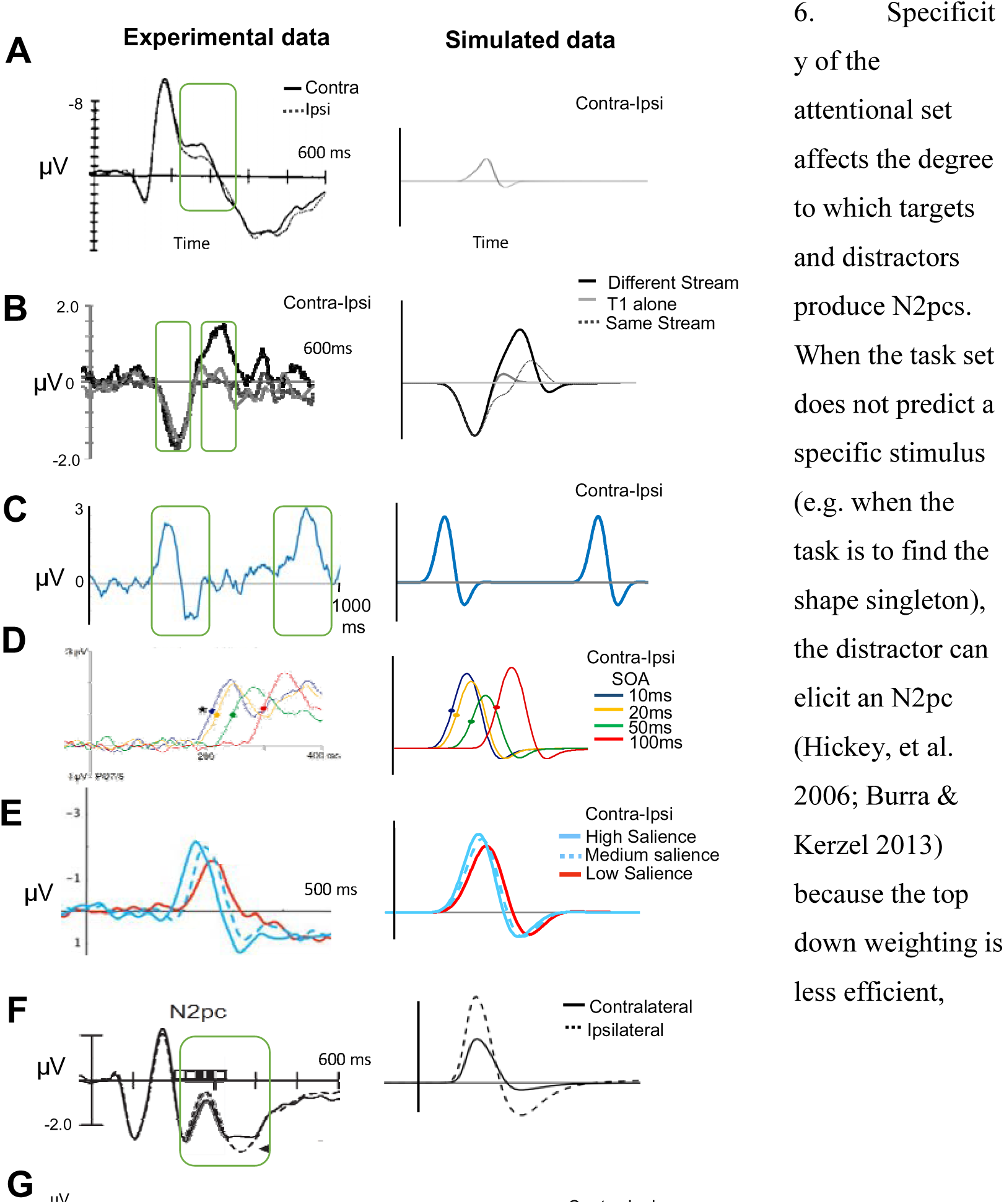
EEG constraints and simulations. Note that polarity is oriented according to the original source and thus switches between panels (a) A laterally presented target causes a brief contralateral negativity, even if it has a long duration (Eimer 1996; green window added to emphasize the time frame of the N2pc). (b) The N2pc to a second target is muted if it occurs very soon after and in the same location as a preceding target (Tan & Wyble 2014). (c) The second N2pc is of normal size if the two targets are far apart in time (Callahan-Flintoft, Chen, & Wyble 2018). (d) When two highly-salient, unmasked targets are presented in rapid sequence at different locations, the N2pc to the second target is not much delayed (Grubert, Eimer 2014). (e) The N2pc is often followed by a deflection in the positive direction, when the target is highly salient (Töllner, et al. 2011). (f) A laterally presented distractor can trigger an N2pc (McDonald et al. 2012). (g) With a highly predictable or salient target, the distractor produces a minimal N2pc and has little effect on the target’s N2pc (Exp 2). When the target set is less specific the distractor has a greater effect on the target N2pc (Exp 1, Hilimire & Corballis 2014).

As a model, RAGNOROC is both an architecture, as well as a specific set of parameters that are calibrated against several decades of data that specify the time course of reflexive attention.

Presumably, this time course reflects an adaptation imposed by other constraints of the visual system. For example, the operation of reflexive attention has to occur within the time span of a visual fixation, while the eye’s position is relatively stationary. During the time window of a single fixation, the representations throughout the visual hieararchy would be roughly in spatiotopic register, making it easy to determine which information is associated with the same object across different maps.

With the model developed and parameterized, the next steps are to use it as a tool to learn more about the visual attention system, and to assert a series of testable predictions that can measure the validity of the model relative to the human system. We begin with a series of lessons that were learned through the model’s development and then proceed to some more specific predictions. These lessons are points that are largely implicit in many findings, theories and models that have come over the years, and are described here to make the implications of such ideas explicit for the benefit of readers.

### Lesson 1. Attention does not draw a clear distinction between targets and distractors

Experimental paradigms in psychology often designate stimuli as targets or distractors and it is tempting to assume that the mind of the participant adopts the same crisp distinction. However, the visual system is presumably maintaining vigilance for all possible kinds of stimuli (e.g. consider whether the participant would react to an unexpected flash of light in an experimental context). To accomplish this feat of general vigilance, even during a highly explicit visual attention experiment, the visual system must evaluate all stimuli to determine which, if any, should be attended. This idea was critical in two-stage models of attention (Treisman & Gelade 1980, Hoffman 1979), which posited explicitly that stimuli had to be evaluated in sequence to determine whether they were targets. RAGNAROC extends this idea to reflexive attention mechanisms such that, within the confines of the attention map, for at least the first two hundred milliseconds of processing, there is no categorical distinction between targets and distractors.

Rather, all stimuli compete, and attention is deployed to the winners, and the losers are suppressed (though priority is biased towards stimuli that bear target-defining attributes). The implications of this idea become more interesting when we think about tasks with multiple targets of varying priority.

### Lesson 2: Visual Attention as a decision process

In RAGNAROC, the lock-on dynamics (including the enhancement at the attended location, the surround suppression and the suppression of the IG neurons) all serve to generate a commitment to attend to one or more locations for at least a brief window of time (roughly 100ms or so).

These bursts of attentional lock-on provide stability to reflexive attention over the time span of typical visual fixations, and allow the entire visual stream to momentarily synchronize representations across the multitude of maps distributed throughout the ventral and dorsal streams. This is one means to address the classic notion of binding (Treisman 1996). Without the extra circuitry of the AM, reflexive attention would be prone to jumping rapidly from one stimulus to another, leading to jumbled and mismatched representations in the various maps of the ventral stream.

Even more interesting, however is that RAGNAROC is able to implement these decisions over many possible spatial configurations. In contrast to more standardized approaches in which decisions occur between a fixed set of discrete alternatives, a lock-on state in the AM could be confined to a single point, spread across multiple points, or be distributed across one or more large regions of indeterminate shape.

### Lesson 3: What does the N2pc/ P_D_ complex reflect?

A typical approach in theoretical work is to assign specific roles to particular EEG components. For example the N2pc is thought to reflect attention evoked by a target in some form, while the P_D_ reflects inhibition evoked by a distractor. However, as we note above, there are cases in which targets elicit a P_D_ component and distractors elicit an N2pc. This modeling approach illustrates why it is important to consider the many-to-one mapping between current sources and ERPs. The neutrality of a scalp potential at a given latency could indicate a period of neural inactivity, but it could also be the case that there are strong underlying dipoles that happen to cancel one another out at that particular moment in time. It is therefore crucial to ultimately understand ERPs at their source. In a similar fashion, there are several ways in which neural activity evoked by a stimulus could lead to a negativity or positivity. For example, RAGNAROC illustrates why the N2pc is often followed by a positive rebound after about 100ms, even though the stimulus stays on the screen (Brisson & Jolicouer 2007). Furthermore, the model explains why this rebound can increase to the point of producing a trailing positivity as target salience is increased (e.g. Tollner et al. 2011) despite there being no specific distractor.

### Lesson 4. Experiment outcomes are a mixture of different trial outcomes

In RAGNAROC, trial-to-trial variability in the simulations accounts for uncontrolled sources of variability (e.g. spontaneous fluctuations in attentional focus on the part of the subject) and is essential for simulating different levels of accuracy. More importantly, the model clarifies how differences in the magnitude of an effect could reflect variation in the frequency of a given outcome, rather than differences in the size of the effect within each trial, a point that was also emphasized by Zehetleitner et al. (2013). For example, a given experiment that exhibits a weak attentional capture effect by a salient distractor, may in fact have a very strong capture effect, but only on a minority of trials. Likewise, a manipulation that produces a stronger N2pc in one condition may be altering the proportion of trials that contain an N2pc rather than the amplitude of the N2pc itself. This occurs with eye movements as well. For example, van Zoest, Donk, & Theeuwes (2004) demonstrated that the proportion of eye movements towards a target vs a distractor varies proportionately across different viewing conditions.

### Lesson 5. Understanding reaction time costs in attentional capture

The term *attentional capture* typically refers to a behavioral phenomenon of slowed responses to a target due to the presence of a distractor, but what exactly causes the reduced performance? In RAGNAROC, there are three possible patterns of attentional allocation when a target and at least one distractor are presented together. First, the target might trigger attention and suppress attention to the distractor(s); second, the target and distractor might trigger attention together; and third a distractor might trigger attention and suppress attention to the target. Each of these three possibilities produces a different RT for the target.

RAGNAROC predicts that RTs would be fastest when the target is attended and the distractor is suppressed because this reduces interference caused by distractor processing. When both the target and at least one distractor are attended (i.e. simultaneous attention), RTs to the target would be slightly slowed because simultaneous lock-on states, while stable, are often slightly smaller compared to a case in which the target is dominant. The final case produces the slowest RTs because the target is not enhanced by attention which reduces the strength of evidence for that target.

RAGNAROC also predicts that any given experimental block of an attentional capture experiment is composed of a combination of these three outcomes, with proportions determined by the relative priority of the targets and distractors. Thus, even in a paradigm that has minimal evidence of attentional capture at the group level, the distractor may nevertheless trigger the deployment of attention on a subset of trials depending on variation in the subject’s attentional focus.

### Lesson 6. Architectural answers to the bottom-up/top-down attentional capture debate

One of the most enduring discussions in the attentional literature is whether bottom-up stimuli are always able to capture attention, or are top-down attentional control signals able to override bottom-up salience. Driving this debate are classic findings that some kinds of distractors elicit capture costs consistently, even though they are never task relevant (Theeuwes 1991). In others studies, capture effects seem to be entirely contingent on top-down settings (Folk, Remington & Johnston 1992). This debate has continued without a clear resolution.

In the model, there is a sense to which bottom-up selection occurs prior to top-down guidance because of the anatomical ordering of early vs later stages of processing. Differences in physical salience are represented at the junction between EV and LV, and differences in task-related attentional set are represented between the LV and AM. This means that a difference in physical salience will often manifest in the AM prior to a difference in task relevance simply because the EV neurons are earlier in the processing hierarchy, which allows them to determine which stimuli in the LV will cross threshold first. Figure 16 compares the time course of activation bumps generated by highly-salient, irrelevant stimuli, to less-salient but task relevant stimuli. Thus, the model exhibits a form of precedence that is in general agreement with Theeuwes Atchley & Kramer (2000). Moreover, this result is not due to specific parameter values, but rather is an outcome of the model’s feedforward architecture. Since salience differences are thought to be processed earlier in the hierarchy (Zhaoping 2002), highly salient stimuli will tend to activate their corresponding LV nodes earlier than less salient stimuli. However, this temporal advantage does not mandate that salient stimuli will always be attended first, since a strong top-down weighting can allow a task-relevant, but lower-salience stimulus to establish a lock-on state more quickly than an irrelevant, higher-salience stimulus.

**Figure 16.**
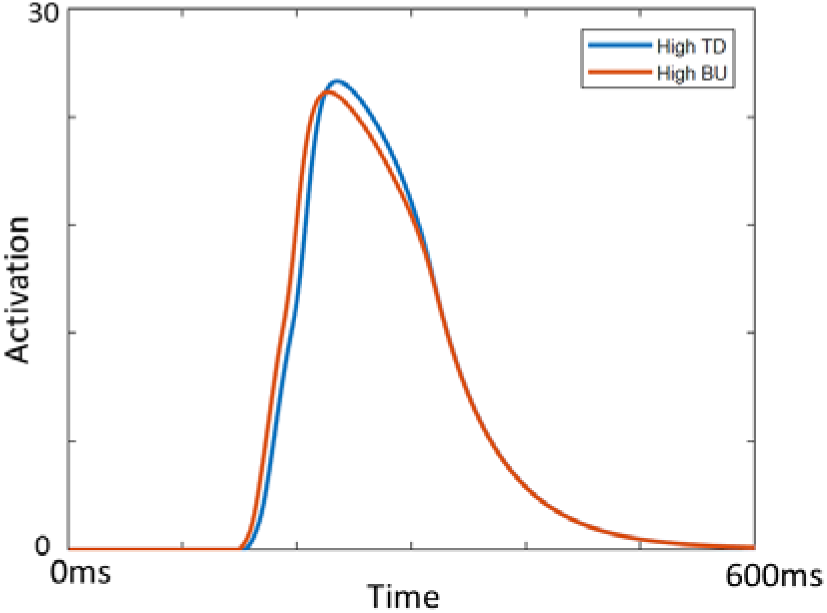
Simulation of the time course of attention map activation for two stimuli that have similar attentional priority, except that one has high salience and a low bottom-up weighting (BU: .2, TD: .15) while the other has the reverse (BU: .15, TD: .2). Despite the higher peak of the high-TD stimulus, the high-BU stimulus has an earlier peak. This is an overlay of two traces; the two stimuli were simulated separately and had the same onset.

### Lesson 7. Architectural answers to the singleton-detection mode debate

Another crucial issue in the attentional capture debate has been the idea that the singleton detection mode allows the system to select unique information for any attribute dimension (e.g. the red item among green items). The advantage of such a mode is that it does not need to be configured in advance for a specific value, preferring equally a red among green items or a green among red items. It has been suggested that subjects use singleton detection when looking for a target that has a unique property, such as a color or form singleton(Bacon & Egeth 1994).

However, a limitation of singleton detection mode is that it cannot be directed towards a specific dimension. Thus, using singleton mode to detect an oddball shape will also prioritize an oddball color.

Models like RAGNAROC make singleton-detection mode straightforward; it is the lack of a strong top-down set, which thereby allows stimuli with high physical salience to dominate the computation of attentional priority. This explains the observation that singleton detection mode cannot be specific for a given dimension. Moreover, since singleton mode is effectively the absence of a top-down set, it is the default search policy (Bacon & Egeth 1994; Lamy & Egeth 2003).

### Lesson 8: Architectural answers to the distractor suppression debate

Competing accounts of inhibitory control in reflexive attention pit the notion of a suppressive surround (Mounts 2000; Cutzu & Tsotsos 2003; Tsotsos 2011) against accounts in which inhibition is selectively deployed to distractor locations (Cepeda et al. 1998; Gaspelin et al. 2015). RAGNAROC illustrates how readily a single model can exhibit both behaviors depending on the paradigm that is being used. A spatial gradient in AM->IG connectivity simulates the surround inhibition effect of Mounts (2000). However, within that surround field, inhibition is selectively applied to the locations of stimuli as a function of their spatiotopic distance to the lock-on state.

RAGNAROC thus explains why the Mounts paradigm and other paradigms which also surround the initial target with distractors such as Cutzu & Tsotsos (2003) were so successful in eliciting the inhibitory surround, while other paradigms have no clear pattern of inhibitory surround (e.g. Wyble & Swan 2015). In the Mounts paradigm, the first target is surrounded by a large number of simultaneously presented distractors. This display is followed immediately by a second display that is used to probe the state of attention. According to RAGNAROC, the dense field of distractors in the first display of Mounts (2000) plays a key role in revealing the shape and size of the inhibitory gradient, since each of those distractors will elicit inhibition in their location, and this inhibition will affect the following target. For paradigms in which the initial target is not surrounded by a dense field of distractors (e.g. Wyble & Swan 2015), the IG neurons in the large area surrounding the target are not stimulated by input from the LV and therefore the only inhibition that is actually expressed in the AM is that immediately surrounding the target’s lock-on state. Figure 17 illustrates a comparison between cases where a target is surrounded by distractors and when it is not.

**Figure 17.**
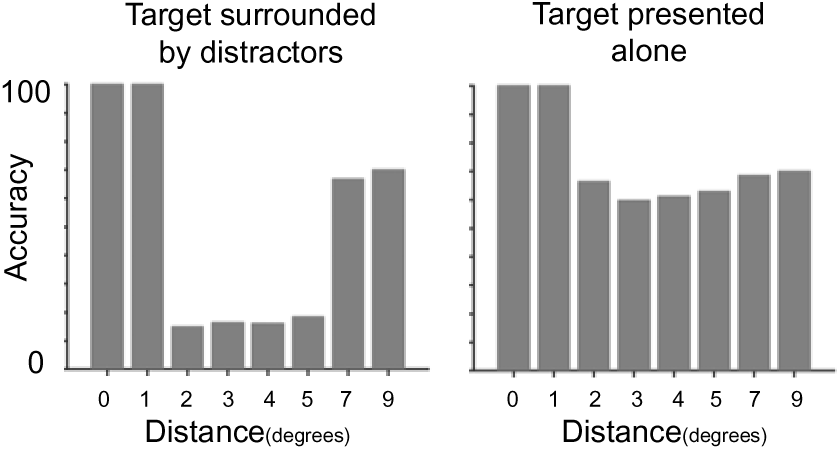
Simulation of the surround inhibition effect of Mounts, showing that the strength of the inhibitory surround is dramatically enhanced by the presence of surround distractors as in the original Mounts (2000) paradigm. A target presented by itself elicits a much weaker inhibitory surround.

### Lesson 9. The Competition for attention can result in a tie

The conventional notion of spatial attention is that it behaves like a spotlight, focusing on only one location at a time. This explanation provides a ready explanation for cueing costs and attentional capture effects, since attention directed at one location can therefore not be at another. However there is also mounting evidence that attention can be deployed simultaneously at two distinct locations (Bay & Wyble 2014; Bichot Cave & Pashler 1999; Kyllingsbaek & Bundesen 2007; Kawahara & Yamada 2012; Goodbourn & Holcombe 2015; see also the possibility of having multiple attention pointers or FINSTs; Pylyshyn & Storm 1988). Of these, Goodbourn & Holcombe provide what is arguably the most compelling evidence of the simultaneity of attentional deployment by measuring the time course of selection at two discrete locations and finding essentially no lag for one vs two simultaneously cued locations. The RAGNAROC model provides an explanation for these seemingly incompatible sets of findings. The circuitry in the attention map elicits a competition for attention between nearly concurrent stimuli, however it is a competition in which there can be multiple winners, which allows simultaneous attention for two stimuli of approximately equal priority.

### Lesson 10. Reflexive attention may have almost unlimited capacity

A common assumption of cognitive theories is that attentional limitations play a key role in determining performance in complex tasks. However, attention is a broad concept and it is often difficult to understand exactly what forms such limits take. In many cases, attention is equated with the ability to “process” information, which includes some mixture of identification, decision making, response generation, and memory encoding.

RAGNAROC embodies a specific definition of attention, which is the reflexive enhancement of feedforward excitation at a given location in the visual field deployed reflexively in response to a stimulus. In the model, this form of attention has no clearly defined limit in terms of the number of attended locations, since the increase in gain could occur at nearly any number of locations. Thus, the model proposes that the earliest stage of attentional selection may operate without strict capacity limits. Of course, subsequent stages of processing are surely limited. For example, even if four stimuli produced simultaneous lock-on states, encoding them all into memory at the same time would produce interference. Parallel selection at early stages does not necessarily entail parallel processing at later stages.

### Lesson 11. Attention can be suppressed without suppressing the representations

It is often suggested that distractors are inhibited during the selection of target information but RAGNAROC elucidates an important distinction between suppressing the representation of a stimulus itself vs suppressing attention at the stimulus’ location. Suppressing a stimulus representation entails direct inhibition of the neurons that represent the attributes and features activated by that particular stimulus (e.g. Reynolds & Heeger 2009; Beuth & Hamker 2011) with the potential to eliminate the active representation of that information from the nervous system. On the other hand, suppressing attention at the location of a stimulus, as in RAGNAROC, preserves the original information of the stimulus at the earliest layers of the visual system.

It is difficult to clearly distinguish between the two implementations of suppression using observations of accuracy or reaction time, since both will reduce the ability to respond to a stimulus. However this difficulty illustrates an advantage of the modeling approach, since models are able to clarify distinctions of implementation that are not otherwise obvious (see also Lu & Dosher 1998 for an illustration of how models of noise exclusion can provide a more specific inference about the mechanisms of attention with the use of psychometric curves).

Moreover, the model illustrates why it would be advantageous to suppress attention, rather than the representation. Suppressing the representation of a stimulus would require an enormous number of long-range connections to deliver inhibition to the appropriate neurons throughout the set of LV maps. Suppression of attention is much simpler to implement, since the inhibitory circuitry is entirely self-contained within the AM.

#### 6: Review of other theories

There is an enormous literature of theories and models of visual attention and the mechanisms in RAGNAROC are inspired by this work. To this base of knowledge, the model contributes the following:

- mechanism for making rapid decisions regarding which spatial locations of the visual field should be selectively suppressed
- explanation for the N2pc/ P_D_ complex as a neural correlate of this decision process
- contact with the empirical literature for behavioral and electrophysiological correlates of reflexive attention^7^

• What follows is a comparison and contrast with other existing models of spatial attention.

### 6.1 Models inspired by neurophysiology

There is a family of models and theories of visual attention inspired by single unit neurophysiology in monkeys. Some of the research in this domain explores the properties of attention in spatial and feature domains. For example, the normalization model of Reynolds & Heeger (2009) proposes that the neural response to any given stimulus is downweighted by the activity of nearby stimuli. Thus, when one stimulus is attended, other stimuli in the vicinity will evoke less activity, all else being equal. The normalization model provides a straightforward, neurally plausible mechanism for the effects of attention at the level of single-unit data. Beuth & Hamker (2015) provide a more detailed account of how attention can be mediated at the level of cortical representations. Such models interface directly with single-unit data from a variety of cortical areas, although they do not explain the decision-making aspect of spatial attention that is the focus of RAGNAROC. Instead, attention is directed by mechanisms that are outside the scope of those models, making them complementary to this model. However, we consider it an open question whether the suppression of attention is best explained with computations at the local circuit level within earlier cortical areas, as in Reynolds & Heeger (2009) and Beuth & Hamker (2015) or at a superordinate level as in RAGNAROC.

Another well-known theory of attention is biased competition, in which stimuli compete with one another for representation in a neural field with overlapping receptive fields. The competition is biased in favor of neurons that respond preferentially to task-relevant information (Moran & Desimone 1985; Desimone & Duncan 1995). There is an important point of correspondence between biased competition (BC) and RAGNAROC, which is that both incorporate an initial period of non-selective processing before the deployment of spatial attention. However, the RAGNAROC and BC models differ in the specific mechanism of attentional enhancement, since BC implements attention as a contraction of receptive fields around the target stimulus while RAGNAROC uses spatially selective multiplicative enhancement. The difference is key because at the core of BC is the idea that representational space is a limited resource, which strongly limits the ability to attend to multiple locations at once. In RAGNAROC, this reflexive form of attention has fewer limits and thus can be deployed to large regions or multiple locations. With that being said, the effect of attention in RAGNAROC appears similar to the single-unit data that inspired the BC model, since an attended stimulus in our model would cause inhibition of attention to nearby stimuli in the attention map. Consequently, activity in LV for an unattended stimulus drops sharply after the initial burst of activity compared to an attended stimulus because the increasing inhibitory feedback from II is not countered by an increase in attentional enhancement. Therefore the model replicates a similar effect in neural activity as BC (Figure 18), but the cause is a suppression of attention, rather than shift in receptive fields.. As in biased competition this suppression would not occur for stimuli that were spaced farther apart since there is a limit to the spatial extent of inhibition in the AM (Figure 18, right panel).

**Figure 18.**
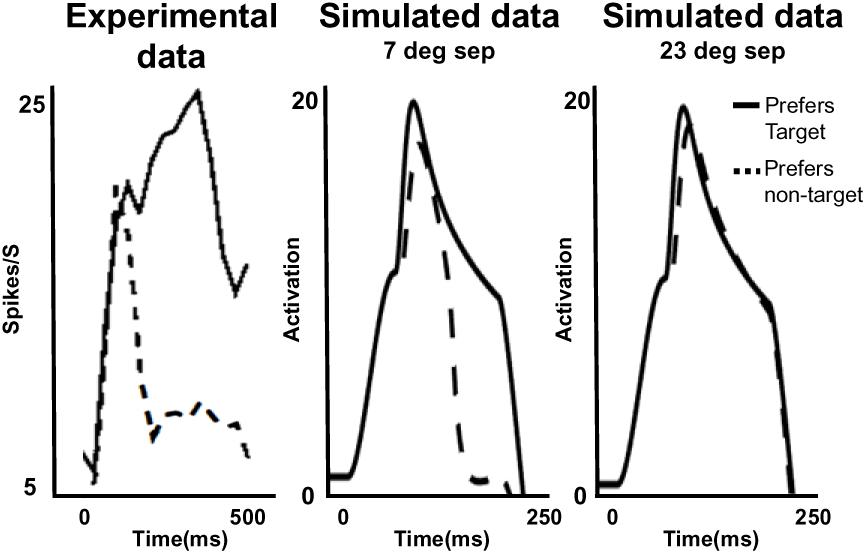
Simulation of the biased competition effects in Desimone & Duncan (1995) in which the activity of a neuron selective for the unattended stimulus of a pair becomes dramatically less responsive to that stimulus after an initial period of processing, but only for stimuli that are spatially proximal. Simulated membrane potential is used as a proxy for spiking rate since RAGNAROC is not a spiking model.

### 6.2 Theory of Visual Attention

The Theory of Visual attention (Bundesen 1990) is a mathematical abstraction of the process of attending to and perceiving one or more stimuli in a single display. In TVA, there are two ways to prioritize certain kinds of information selection: *filtering*, and *pigeonholing*. *Filtering* involves upweighting the priority for certain features, which increases the rate at which stimuli possessing those features attract attention. This is similar to attentional control settings in RAGNAROC. The *pigeonholing* mechanism relates to how efficiently certain kinds of information are categorized, which allows them to be reported and remembered. The TVA model thus represents two distinct types of attentional control setting, which might also be described as *key feature* and *response feature* (Botella, Barriopedro, & Suero 2001). RAGNAROC differs from TVA in that it provides a more complete model of the neural mechanisms associated with the computation and use of priority to direct spatial attention. The TVA model, on the other hand, provides a concise mathematical formulation of how two different kinds of filters interact to facilitate perception. A neural implementation of TVA has been proposed (Bundesen et al. 2011), however it is less clear how such a model would scale up to a full working specification, since it requires a large scale winner-take-all implementation to complete attentional selection, which is likely to be incompatible with white matter constraints. Given the similar role of priority within the two models, it could be fruitful to consider RAGNAROC as a neurophysiologically plausible front-end for computing the priority-based competitive selection process and the TVA as a specification of subsequent processing.

### 6.3 Guided Search

. To better understand the complexites of the visual search literature, the Guided Search model (GS; Wolfe 1994) simulates how top down goals interact with bottom up salience signals to determine likely target locations. Like RAGNAROC, this model attempts to explain how the visual system mediates the balance between salience and task relevance. Its focus is on a longer time scale than reflexive attention, and incorporates both overt and covert forms of attention. RAGNAROC is complementary to this model, by explaining attentional dynamics at a short time scale, and with a greater emphasis on decisions, selective inhibition and neural correlates.

### 6.4 Feature Map Models

Another class of models simulates spatial attentional effects across sheets of neurons corresponding to different visual features. Perhaps the most canonical of such models is the salience model of Koch & Ulman (1985), which is architecturally similar to RAGNAROC. A descendent of this model is often invoked as a benchmark in computer vision algorithms (Itti, Koch & Niebur 1998).

In such models, feature detectors in multiple channels (e.g. luminance, color, motion flicker) project to a master salience map that ultimately makes decisions about where attention will be deployed using a simple winner take-all mechanism, coupled with a form of memory that erases salience values at recently-visited locations. Like RAGNAROC, this model uses salience as a common currency across all stimuli in the visual field, and would be able to simulate capture effects. The Itti, Koch & Niebur (1998) model has been foundational in understanding how simple mechanisms can reproduce complex gaze behavior when iterated over many distinct feature dimensions and levels of scale. Also, because the Itti et al. model simulates responses to pixelwise visual data, and can be compared against visual fixation data from human subjects, it set the stage for a generation of computer vision research.

Itti et al.(1998) and RAGNAROC address phenomena at different time scales. The former is intended as a model of gaze behavior on time scales of a second or more, involving multiple fixations. RAGNAROC is developed to understand how covert attention deployment is computed anew with each visual fixation or significant change to the visual display. Moreover, the salience map in Itti et al.(1998) lacks the decision mechanisms to suppress distractors without first generating an overt gaze response to the distractor. The two models are thus complementary; they operate at distinct time scales, emphasize different kinds of processes and simulate fundamentally different kinds of data.

Other models provide more direct simulations of the neural processes of enhancement in neural sheets. The Selective Tuning model by Tsotsos (1995; 2011) implements a form of inhibition in which detection at an upper level of the hierarchy produces surround inhibition at earlier layers of the hierarchy. This model is perhaps the most well-formulated attention model that has ever been proposed since it proposes a gating control circuitry that allows information to be effectively linked across differences in spatial invariance. Selective Tuning would reach several of the benchmarks described here, but does not propose a means to selectively inhibit distractors. It applies inhibition in a region surrounding a target, irrespective of the presence of distractors.

Moreover, decisions to deploy attention are made independently for different stimulus dimensions and it is not precisely formulated how cross-dimensional competition between stimuli would be implemented at the time scale of reflexive attention (see p121, Tsotsos 2011).

### 6.5 Resonance models

Another variety of models uses excitatory resonance to determine the boundaries of a region that should be attended. Amari (1971) described a general framework for simulating neural field dynamics in densely interconnected networks having a single layer. These ideas have been adapted to simulations of visual attention by assuming that the neural field has shared spatial topography with the visual system. For example the attentional shroud (Fazl, Grossberg & Mingolla 2009), is a means to delineate the boundaries of an object and then ensure consistent focus on that object during learning. Similarly, models such as MORSEL (Mozer & Behrmann 1990) and SAIM (Heinke & Humphreys 2003; Heinke, Humphreys & Tweed 2006) use iterative processing between interconnected neurons to discover the region that satisfies competing constraints from a variety of sources, such as visual input, templates and top-down goals. Other examples include Zirnsak, Beuth & Hamker (2011) that simulate the temporal dynamics of attentional competition in response to one or more stimuli; and Lanyon & Denham (2004) which simulate eyegaze during visual search as a product of interacting attentional systems.

These resonance theories are similar to the lock-on states, which also emerge through iterative processing of interconnected, topographic neural fields. What RAGNAROC contributes to these models, apart from the more explicit link to EEG components, is a mechanism for selective suppression that complements the mechanism for selective enhancement.

## 7. Predictions

RAGNAROC is part of an ongoing investigation that involves a cyclic iteration between theory and experiment. Driving this cycle are *a-priori* predictions, that provide a roadmap for future experimental work to diagnose the model’s validity. By publishing these predictions in advance of testing them, we minimize the file drawer problem, which occurs when model tests are selected for publication after the results are known. Furthermore, our goal here is to specify an ambitious set of predictions, with the expectation that some of them should be inaccurate. Since all models, being abstractions of the real system, are wrong by definition (Box 1976), the prediction/testing cycle should be most efficient when there is a mix of true and false predictions. True predictions give evidence that the model has at least some resemblance to the underlying system. However, it is the false predictions that are truly valuable, for they indicate where the model is inaccurate, and thereby guide further development of the theory. These predictions are divided into three categories below.

### 7.1 Competition within the attention map

These predictions concern the essential architecture of the model. Failure to validate them would require at a minimum, significant parameter or architectural changes. In RAGNAROC, the competition for attention exists between all stimuli, and the priority values of the stimuli are the common currency with which they compete. Since the attention map does not represent the distinction between targets and distractors, the following predictions should obtain:

#### Prediction 1. Lower priority targets will elicit AM suppression

In RAGNAROC, input to the attention map does not distinguish between targets and distractors. A counterintuitive prediction of this assumption is that when a display contains two targets with sufficiently different priority values, the lower priority target will lose the competition and be treated as a distractor by the very first pass of reflexive attention. This would mean that the weaker target will elicit a weak N2pc when presented laterally, followed by a clear P_D_ component as if it had been a distractor. In terms of behavior, the location of the low-priority target should exhibit the same lower probability of probe letter reporting as the salient distractors of Gaspelin et al. (2015). If the high-priority target were omitted from the display in other trials, the weaker target would now elicit an N2pc and increase, rather than decrease behavioral responses to a probe at its location. Target priority could be manipulated either by varying the salience of targets or their proximity to the task-defined attentional set in some feature dimension, such as color (Becker, Folk & Remington 2013).

#### Prediction 2. Higher priority distractors will more often elicit a lock-on state

A similar kind of prediction can be made for distractors of varying priority. If a display consists of only distractors of three or more clearly discernable levels of salience (e.g. by adjusting their relative luminance), the distractors will elicit N2pc and P_D_ components as if the most salient distractor were a target and the next most salient distractor were the key distractor in the additional singleton paradigm. The most salient distractor will also capture attention resulting in improved accuracy and reduced reaction times for probes (e.g. Gaspelin et al. 2015) at its location. Conversely, probes at the second-most salient distractor location will be less well perceived than distractors at the location of the least salient distractor. This prediction stems from the fact that the amount of inhibition delivered to the location of a lower-priority stimulus in the AM is proportional to its priority. Testing this prediction would require embedding distractor-only trials within a larger set of trials that contain targets as well. Some of these trials would contain probe letters as in Gaspelin et al. (2015)

Note that that there is conflicting evidence about the ability of distractors to elicit an N2pc. Distractors that are highly salient on a different dimension than the target (e.g. color singleton distractors with shape singleton targets) elicit an N2pc, while a difference in salience along the same dimension (color) does not (Gaspar & McDonald 2014).

#### Prediction 3. PD amplitude is modulated by spatiotopic distance between target and distractor

The selective inhibition of distractors in the AM is driven by a pattern of localized excitatory connectivity between AM neurons and IG neurons. This means that competition between different stimuli tapers off at large inter-stimulus distances. Thus, all else being equal, when a salient distractor is placed very far from a target, the PD will be reduced in size relative to when the target is proximal. It is difficult to make a specific prediction about the spatial scale of the falloff without more data to constrain the model at this point, but it should be on the order of 10 degrees or more according to the spatial extent of LAI as found in Mounts (2000).

### 7.2 Unified Attentional Map

A central theme of the RAGNAROC architecture is that the competition for reflexive attention is confined to a small region of neural tissue this is sensitive only to stimulus priority. This allows the entirety of the visual system to participate in scene analysis, and yet make rapid, efficient and stable decisions about the allocation of attention. The attention map allows the priority signals generated by different stimuli to compete, taking into account their salience, task relevance, emotional/reward history, or any other potential factor that influences how a given stimulus should be prioritized. This idea of a single, superordinate attention map is shared by many models of visual attention (Itti Koch et al 1998; Zelinsky 2008) but not others (Tsotsos 2011).

#### Prediction 4. Salient Distractors can sustain an existing lock on state

One of the most counterintuitive predictions of RAGNAROC is that once an attentional lock-on state has been established by a target, it can be sustained by a distractor because the attention map is agnostic about target/distractor categories. Attentional control settings bias attention towards the target, but distractors also have excitatory connections to the AM; they just have reduced priority. Thus, in a similar manner as two sequential targets can maintain a lock-on state (Tan & Wyble 2015), a target followed by a distractor should also maintain the lock-on state.

The prediction can be tested by presenting either three targets in sequence (i.e. letters among digits), at an SOA of about 120ms, or two targets separated by a single distractor that is similar to other distractors (i.e. a black digit), or two targets separated by a highly salient distractor (i.e. a red digit). It should be observed that for three targets in a row, the second and third targets elicit small-amplitude N2pcs that peak early (roughly 30ms earlier than the relative latency of the T1’s N2pc). If the middle of the three targets is replaced by a highly salient distractor, the last target’s N2pc should still be early and small in amplitude. However in the case of two targets separated by a non-salient distractor, that last target should evoke an N2pc of normal amplitude and latency since the lock-on state will have partially dissipated during the 240ms lag between the onset of the two targets. In behavior, the salient intervening distractor should result in more accurate report of the following target relative to the non-salient distractor condition, since the highly salient distractor sustains the lock-on state across the temporal gap between the targets.

#### Prediction 5: EEG correlates of lock-on occur regardless of stimulus type

A core finding of the lock-on state presents a straightforward means to test this architectural prediction. In Tan & Wyble(2015), it was found that two targets in the same location produced only an N2pc to the first target, which RAGNAROC explains as a carryover of the attentional lock-on state from one target to the next in the attention map.

However, in that study, both targets were of the same kind (letters among digit distractors). If there is a single attention map, the carryover of lock-on from one stimulus to the next should occur even when T1 and T2 are of different types. Callahan-Flintoft, Chen, & Wyble (2018) provided support for this prediction already by showing that targets could be defined by combinations of shape or color without disrupting the lock-on effect. It is nevertheless possible that the prediction may be falsified if the two targets were even more distinct. For example, RAGNAROC predicts that even if subjects are simultaneously looking for letters and faces of a particular gender, then two sequential targets (either letter-face or face-letter) should produce a clear N2pc only for the first of the two targets. Letters and faces should provide a strong test for the hypothesis since previous work has suggested that they are processed through sufficiently distinct channels in the visual system that the attentional blink evoked by a digit T1 has little effect on a face T2 (Awh et al. 2004). A failure to confirm this prediction would suggest that there are subdivisions of the attention map for stimuli that are highly distinct.

### 7.3 Lock-on states in visual cueing

The RAGNAROC model implements a reflexive form of attention that should be common across many paradigms, including visual cueing. Thus, we should be able to predict behavioral and ERP effects for cueing experiments as well. Ansorge, Kiss, Worschech & Eimer (2011) have demonstrated that cues evoke clear N2pcs at moderate cue-target SOAs (200ms), as we would expect. However shorter Cue-Target SOAs should reveal correlates of the lock-on state.

#### Prediction 6. Lock-on states in visual cueing, valid trials

RAGNAROC predicts that a lock-on state is sustained from one stimulus to the next if they are at the same location and the SOA is on the order of 100ms. Thus, from a behavioral perspective, RAGNAROC explains cueing benefits at short SOAs, if one assumes that a cue initiates a lock-on state that carries forward in time to enhance the target. The model also generates EEG predictions for cueing experiments. Since the N2pc is caused by the formation of a new lock on state, then a validly cued trial with a short SOA (i.e. 100ms or less) should result in a N2pc appearing for the cue, and a dwarfed, early N2pc appearing for the target (c.f. Callahan-Flintoft, Chen & Wyble 2018).

At longer SOAs (e.g. 500ms or more) between the cue and target, the lock-on state elicited by the cue would have disintegrated before the target appeared, with the result that both the cue and the target would produce a typical N2pc.

#### Prediction 7. Spatial separation modulates selective suppression

When a cue and target are not in the same location, then the cue and target will each elicit an N2pc at all SOAs, since the lock-on state elicited by a cue is spatially specific. Thus, a target in an invalid trial always needs to build a new lock on state, which elicits a new N2pc. A failure to confirm these predictions would undercut the applicability of RAGNAROC’s simulation of attention related EEG components to cueing studies and suggest that there are unappreciated differences between the way that targets and cues are processed by the reflexive attentional system.

### 7.4 Lock-On to naturalistic stimuli

#### Prediction 8. Regional deployment of attention

The collective understanding of reflexive attention, as reviewed here, stems almost entirely from studies that use discrete stimuli, such as shapes, characters, images or sketches. While these methods are ideal for experimental control, it is important to apply our theories to stimuli that more closely resemble what the visual system typically encounters, which can be approximated with digital video. Given some assumptions about the computation of salience, RAGNAROC can predict how the spatial and temporal distribution of reflexive attention responds to the physical salience of a series of video frames. Given a time series of salience outputs from a model such as Graph-Based Visual Salience (Harel, Koch & Perona, 2007) RAGNAROC predicts that the competitive inhibitory circuitry of the AM allows attention to flow dynamically across large regions to fill-in gaps in the interior of an object (Figure 19), in a similar manner as *attention shrouds* (e.g. Fazl et al 2009). This is consistent with what would be construed as object-based attention (Duncan 1984) in that the spatial scope of reflexive attention extends to the boundaries of a clearly defined object and also fills in the interior area, even if that interior is not particularly salient. Moreover, the amplitude of activity at various points in the shroud will be relatively constant, which means that attention would be distributed equally across the entire attended area rather than remaining focused at highly salient points. This uniform distribution across an attended region is due to several factors: 1. lock-on states rise rapidly to a ceiling (see Figure 7), 2. the tiled competitive inhibitory circuits within the AM allows lock-on states to exist at many locations simultaneously, and 3. the large receptive fields of the AM encourages causes attention to spread within a salient region..

**Figure 19.**
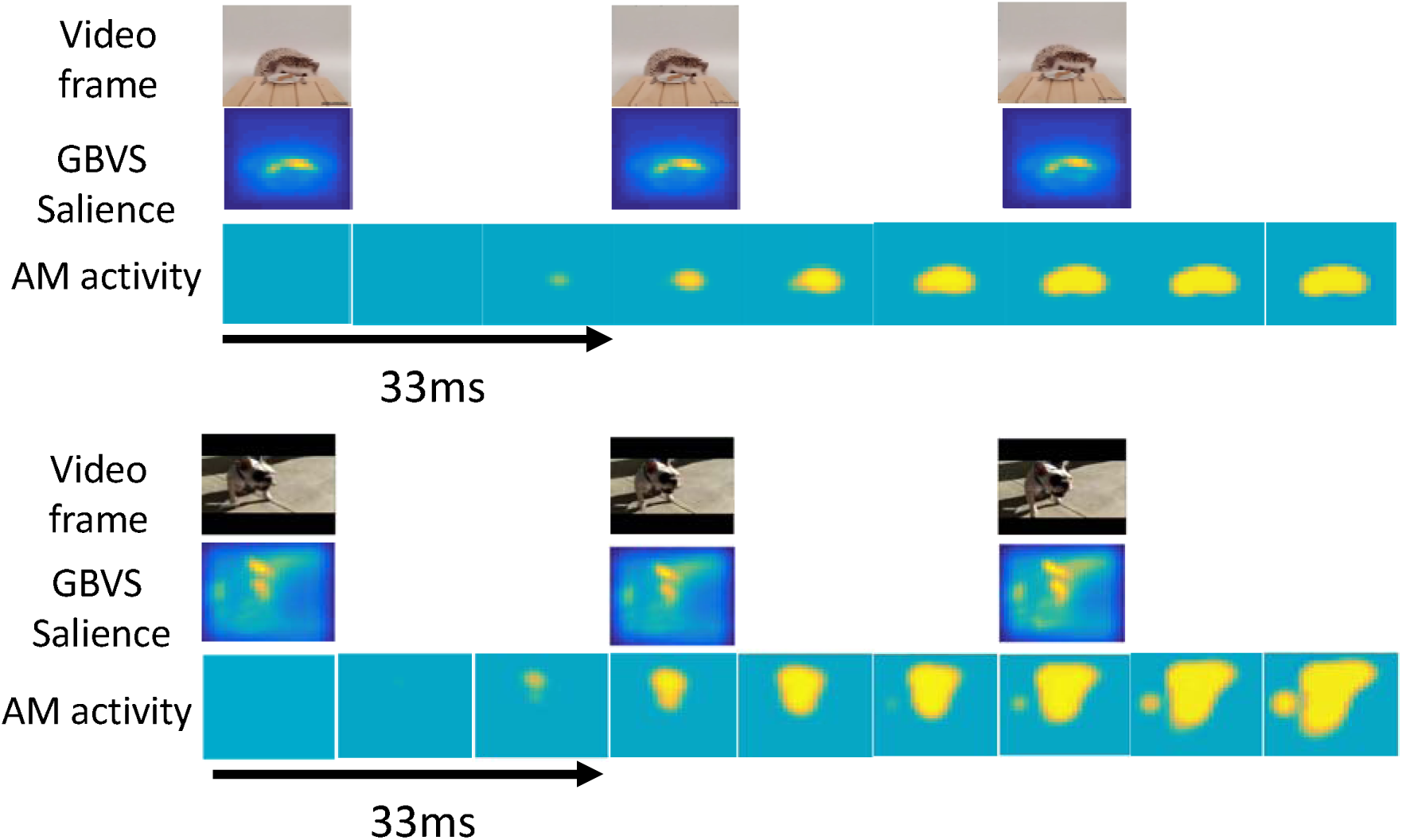
Simulation of AM activity across 99ms of time in response to salience computed by GBVS for three video frames for two different movies. Simulation results for longer videos are provided at the OSF repository for the model.

The key point to take from these simulations is that while attentional cueing and capture effects are typically construed as being anchored to a specific locus and then released to move to the next stimulus (Duncan Ward & Shapiro 1994), the more typical behavior of reflexive attention may be to attend to large regions of space in patterns that shift rapidly to track changes in a dynamic input. The simulations that produce these dynamic shrouds in Figure 19 use the same fixed parameters as the rest of the simulations reported here^8^. Thus, while cognitive experiments demonstrate attentional deployment to discrete locations when stimuli appear and disappear abruptly, these simulations from RAGNAROC show that these effects may be a consequence of the kinds of stimuli that are used in experiments. As we move forward in our understanding of the mechanisms and functions of visuospatial attention it is imperative that we shift our empirical and theoretical focii towards stimuli that have more realistic temporal dynamics.

In this discussion, it is important to emphasize that reflexive attention is only a fragment of the visual attention system, and RAGNAROC lacks, for example, higher-level mechanisms for temporal segmentation that would cause an attentional blink (Raymond Shapiro & Arnell 1992), or explicit object-based attention mechanisms that are driven by border ownership, Gestalt properties and other higher-order visual statistics. Moreover, RAGNAROC is not building event models, meaningful interpretations or computing inference. These other mechanisms would further augment the ability to segment incoming information in a way that allows meaningful representations to be extracted.

To generate this prediction, salience maps were computed for short sequences of frames from two 30 frame-per-second movies from the Moments in Time database (Monfort et al. 2019) using Graph-Based-Visual-Salience (Harel, Koch & Perona, 2007)^9^. These salience maps were provided as input to one LV of RAGNAROC to simulate how attention would evolve over time given such input. The salience input was updated every 33 ms, but the model’s activation was computed at each millisecond.

This is a prediction for which there is currently no clear path to test experimentally and it is provided to inspire the development of new methods. Overt attentional models can easily be tested by tracking eye movements, but here the prediction is of large scale shifts in the shape of covert attention. Probe tasks, such as that of Cepeda et al. (1998) and Gaspelin et al (2015) may be able to chart the evolution of such dynamic covert attentional dynamics.

## 8.0 Conclusions and future directions

Reflexive visual attention is a cornerstone of our visual system’s ability to meet the challenge of rapidly choosing which information to selectively process. A variety of experimental paradigms have provided a wealth of data that we have distilled into a common architecture for controlling the selection and suppression of information. The goal of the RAGNAROC model is to build a theoretical bridge between different paradigms (e.g. visual cueing and capture), and also between different kinds of data (e.g. behavior and EEG). While designing the model to hit its empirical benchmarks, we have developed circuits that use competing attractor states to briefly stabilize the deployment of attention, and to selective inhibit attention at the location of distracting information.

Moving forward, the model’s predictions are intended as a roadmap for further empirical investigation of reflexive attention and for creating links across paradigmatic boundaries. Testing these predictions will provide diagnostic data regarding the model’s validity, but more importantly, will drive further development of the model. For example, it would be useful to provide a more rigorous account of the specific regions of the brain that are responsible for generating these ERPs so as to match not just the time course, but also the scalp topography of the N2pc/ P_D_ complex.

While RAGNAROC is intended as a model of reflexive attention that can be deployed covertly, future work could extend these mechanisms as a partial explanation of the time course of eye movements in visual displays. Doing this would require an additional set of assumptions regarding how activity in the attention map drives the decision to commit visual saccades.

Recent work that explores the time course and spatial distribution of initial saccades in visual search paradigms (e.g. Gaspelin, Leonard & Luck 2017) indicates that initial saccades are directed towards the location of salient distractors when the distractor’s location is not suppressed, but are directed away from salient distractors when that location is suppressed.

These findings suggest that activity in the attention map contributes to the initial decision of where to commit an overt attentional response.

The neural attractor mechanism of RAGNAROC could be incorporated as a front-end onto models of higher order cognitive phenomena. For example, in models of the attentional blink, the time course of target processing is often the central question, and such models have little to say about the time course of reflexive attention. Combining models such as RAGNAROC with models of the attentional blink (e.g. Olivers, & Meeter 2008; Wyble Bowman & Nieuwenstein 2009; Taatgen, et al. 2009) has the potential for expanding our understanding of the spatial and temporal dynamics of attention out to the order of multiple seconds.

Another crucial area of application will be to simulate how attention is deployed to stimuli that change in real-time, such as movies. Presumably, the time course of attentional engagement and disengagement that are simulated here has relevance to the typical temporal dynamics of movement by objects and people in the kinds of settings that are informative for human vision.

## Acknowledgements

This work was performed with the support of NSF grant 1734220 to B. W. and the National Natural Science Foundation of China grant 31771201 to H.C. Author B.W was responsible for the writing. C.C.F did most of the modelling with assistance from B.W. Author T.M. developed the non Matlab version of the model for dissemination on Jupyterlab with assistance from B.W. and Garrett Sullivan. Authors H.C. and H.B. were responsible for conceptual input and advice. The authors also wish to thank, in particular order, Dirk Kerzel, Alex Holcombe, and Nick Gaspelin, for helpful comments.

## Appendix

MATLAB Code for running the simulations is available on the OSF at https://osf.io/rwynp/ Appendix 1, Equations

In simulation, these equations are simulated using Euler integration with time steps of 1ms. A floor function (implemented here as Max in eqs 1.14 and 1.21 is used to prevent currents from going negative, which adds stability to the simulation at discrete time steps. The Max function is also used in equation 1.15 to ensure that attention modulation of the EV->LV pathway can only be excitatory. The Max function also plays a role in limiting the input from the two sources to the IG neurons to ensure that neither the AM, nor the LV excitation is sufficient to allow them to cross threshold (in other words, a dendritic AND gate (Shepherd & Brayton 1983). These uses of Max functions are not intended as a statement of biological plausibility of synaptic processing, but rather as simplifications to permit a slightly simpler architecture and coarser time steps.

## Early Visual layer

These are the activation equations for each neuron in the EV layer, and note we represent each possible stimulus (T1 and T2) as having distinct EV neurons to reflect the fact that two distinct stimuli will activate distinct groups of neurons in V1 even if presented at the same location. These equations match those specified by O’Reilly & Munakata (2001) where ***Input***, represents the presence of an input stimulus at a given location and time point (either 1 or 0) and ***EV*** represents the activation level of that neuron. ***dtVM*** is a time constant that dictates the rate of change of a neuron by scaling the excitatory, and leak currents. ***EE*** and ***EL*** are the reversal potentials for excitatory and leak currents.

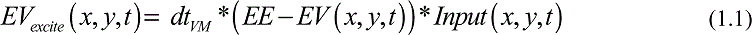

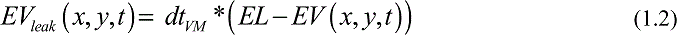

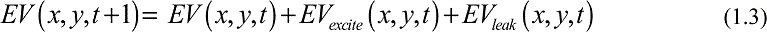

## Late Visual layer

The LV neurons have essentially the same dynamics except that they receive input from a region of EV neurons and the value of the input is scaled by a square-masked Gaussian profile, (***GRF***) for computational efficiency.

The variable ***Attn*** is the current value of attention as determined by activity at the corresponding location in the AM. ***EI*** is the reversal potential of the inhibitory current. ***IItoIT*** is a parameter that determines the strength of the feedback inhibition interneurons for each neuron. ***BU_type_*** is a parameter that reflects the physical salience of a given stimulus type.

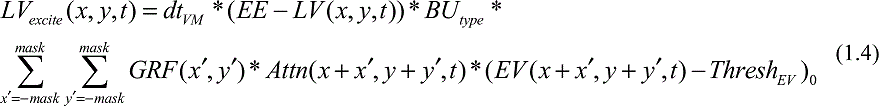

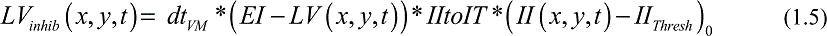

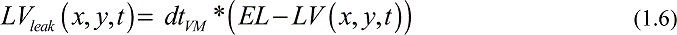

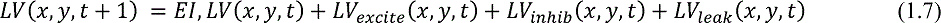

The II neurons govern the feedback inhibition of the LV neurons following a similar dynamic as the EV.

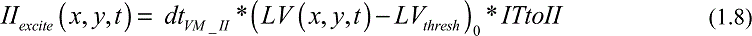

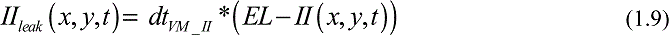

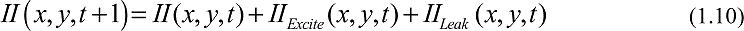

## Attention Map

The AM neurons receive input from all LV maps (1-*n*) scaled by the same masked Gaussian profile ***GRF***. ***LAI*** is a parameter that controls the magnitude of inhibitory suppression from the IG to the AM neurons. ***TD_type_*** is a parameter that determines the top-down task relevance for a given stimulus.

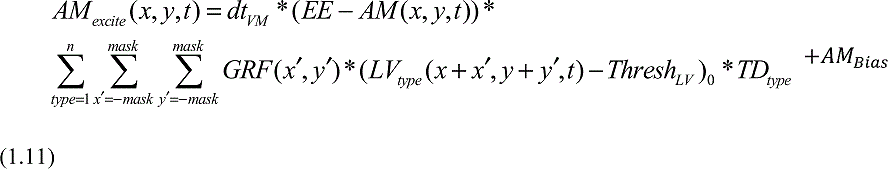

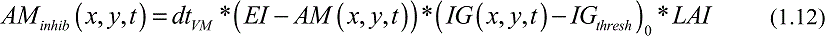

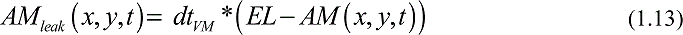

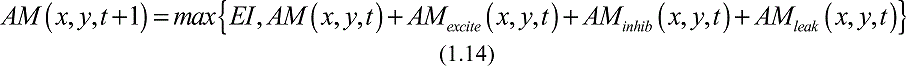

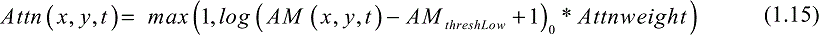

The IG neurons within the Attention Map receive joint input from the LV and the AM. For each IG neuron, the input from each of those two sources has a ceiling value (***MaxInputtoIG***). Thus, the input from the LV and the AM to each IG neuron is computed separately.

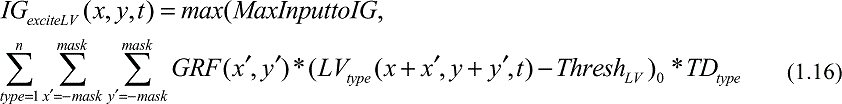

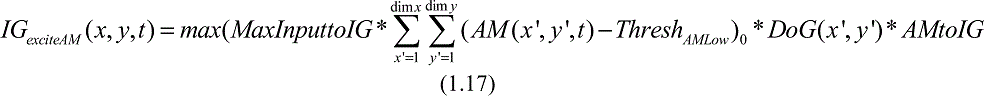

*DoG* is the difference of two Gaussians as specified below.

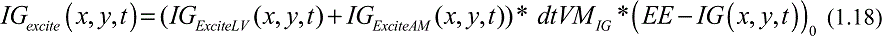

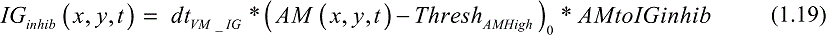

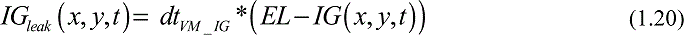

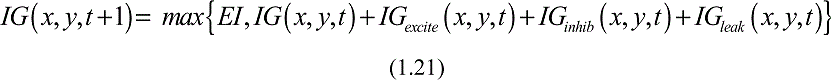

Gaussian Profile

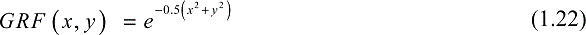

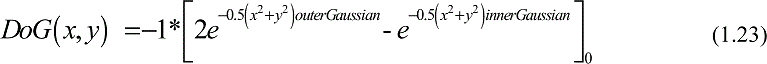

EEG Scalp Voltage

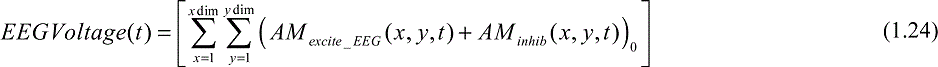

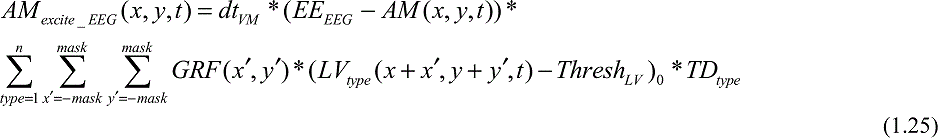

Fixed Parameters:

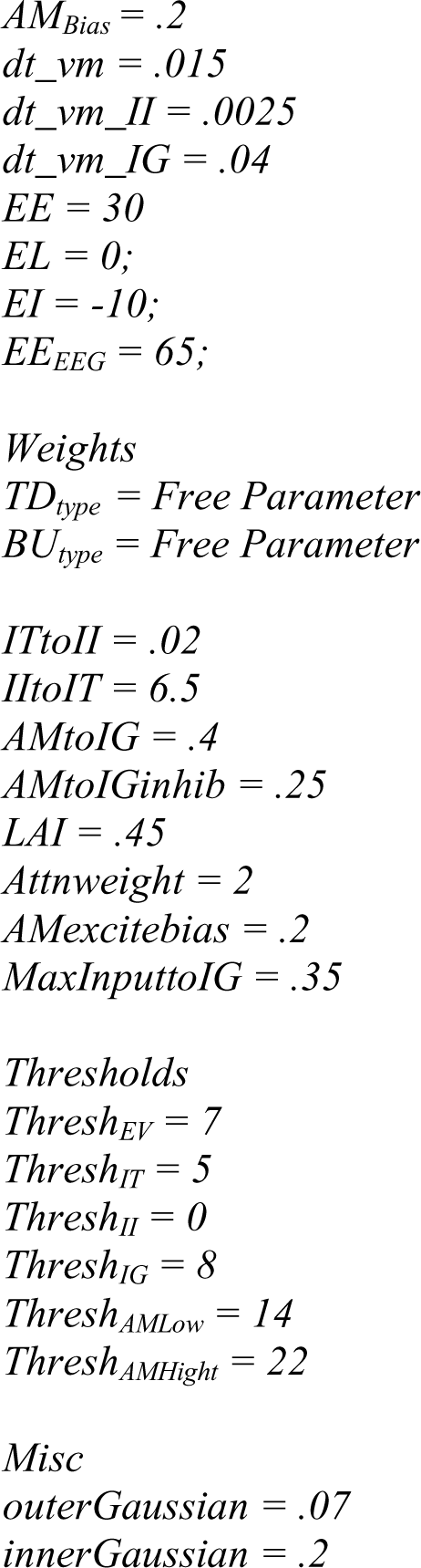

Appendix 2, Fitted Parameters

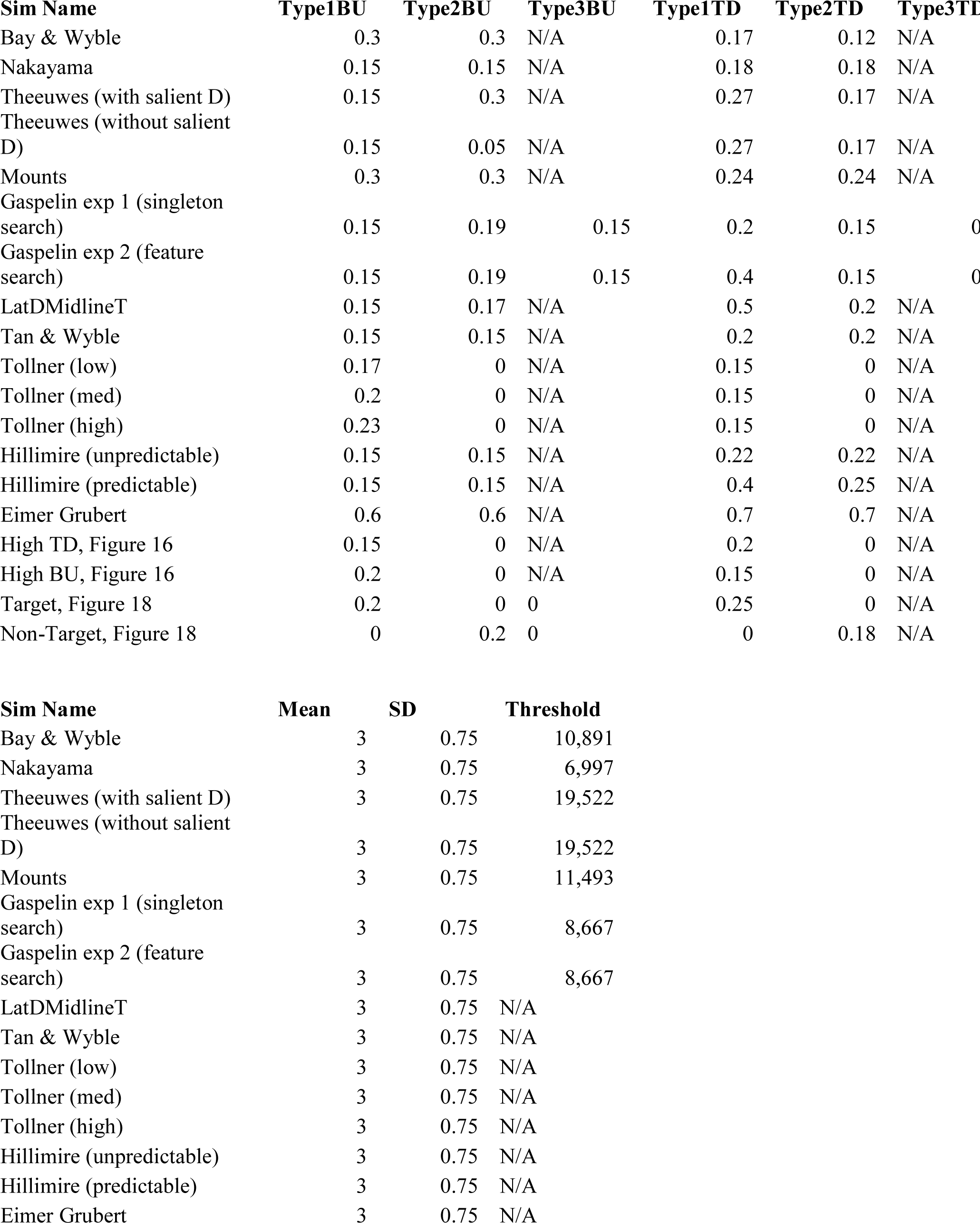
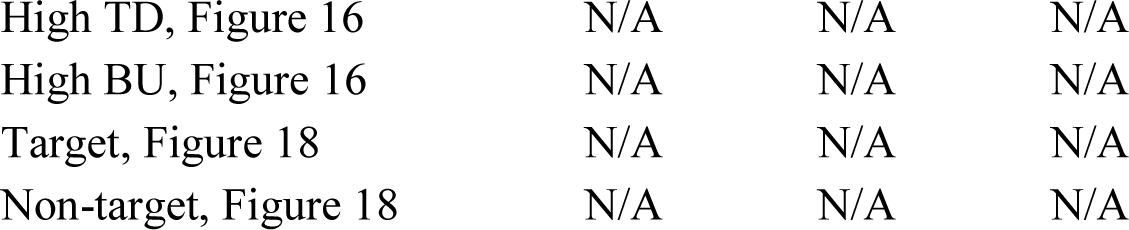

| **Sim Name** | **Type1BU** | **Type2BU** | **Type3BU** | **Type1TD** | **Type2TD** | **Type3TD** |
| --- | --- | --- | --- | --- | --- | --- |
| Bay & Wyble | 0.3 | 0.3 | N/A | 0.17 | 0.12 | N/A |
| Nakayama | 0.15 | 0.15 | N/A | 0.18 | 0.18 | N/A |
| Theeuwes (with salient D) | 0.15 | 0.3 | N/A | 0.27 | 0.17 | N/A |
| Theeuwes (without salient D) | 0.15 | 0.05 | N/A | 0.27 | 0.17 | N/A |
| Mounts | 0.3 | 0.3 | N/A | 0.24 | 0.24 | N/A |
| Gaspelin exp 1 (singleton search) | 0.15 | 0.19 | 0.15 | 0.2 | 0.15 | 0 |
| Gaspelin exp 2 (feature search) | 0.15 | 0.19 | 0.15 | 0.4 | 0.15 | 0 |
| LatDMidlineT | 0.15 | 0.17 | N/A | 0.5 | 0.2 | N/A |
| Tan & Wyble | 0.15 | 0.15 | N/A | 0.2 | 0.2 | N/A |
| Tollner (low) | 0.17 | 0 | N/A | 0.15 | 0 | N/A |
| Tollner (med) | 0.2 | 0 | N/A | 0.15 | 0 | N/A |
| Tollner (high) | 0.23 | 0 | N/A | 0.15 | 0 | N/A |
| Hillmire (unpredictable) | 0.15 | 0.15 | N/A | 0.22 | 0.22 | N/A |
| Hillmire (predictable) | 0.15 | 0.15 | N/A | 0.4 | 0.25 | N/A |
| Eimer Grubert | 0.6 | 0.6 | N/A | 0.7 | 0.7 | N/A |
| High TD, Figure 16 | 0.15 | 0 | N/A | 0.2 | 0 | N/A |
| High BU, Figure 16 | 0.2 | 0 | N/A | 0.15 | 0 | N/A |
| Target, Figure 18 | 0.2 | 0 | 0 | 0.25 | 0 | N/A |
| Non-Target, Figure 18 | 0 | 0.2 | 0 | 0 | 0.18 | N/A |

| **Sim Name** | **Mean** | **SD** | **Threshold** |
| --- | --- | --- | --- |
| Bay & Wyble | 3 | 0.75 | 10,891 |
| Nakayama | 3 | 0.75 | 6,997 |
| Theeuwes (with salient D) | 3 | 0.75 | 19,522 |
| Theeuwes (without salient D) | 3 | 0.75 | 19,522 |
| Mounts | 3 | 0.75 | 11,493 |
| Gaspelin exp 1 (singleton search) | 3 | 0.75 | 8,667 |
| Gaspelin exp 2 (feature search) | 3 | 0.75 | 8,667 |
| LatDMidlineT | 3 | 0.75 | N/A |
| Tan & Wyble | 3 | 0.75 | N/A |
| Tollner (low) | 3 | 0.75 | N/A |
| Tollner (med) | 3 | 0.75 | N/A |
| Tollner (high) | 3 | 0.75 | N/A |
| Hillmire (unpredictable) | 3 | 0.75 | N/A |
| Hillmire (predictable) | 3 | 0.75 | N/A |
| Eimer Grubert | 3 | 0.75 | N/A |

|  |  |  |  |
| --- | --- | --- | --- |
| High TD, Figure 16 | N/A | N/A | N/A |
| High BU, Figure 16 | N/A | N/A | N/A |
| Target, Figure 18 | N/A | N/A | N/A |
| Non-target, Figure 18 | N/A | N/A | N/A |

## Footnote

There is an ongoing debate concerning the ability of top-down goal settings to mediate attentional capture (Awh, Belopolsky & Theeuwes 2012; Failing & Theeuwes 2018), with positions ranging from attention being entirely driven by Top-down factors, to the opposite extreme in which the first stage of attention is entirely driven by physical characteristics of stimuli.

CTSOAs were only evaluated in the range of 100-260ms.

For simplicity we assume that there is only a single cortical area that computes attentional priority, although the functionality would be essentially similar if there is a small family of interconnected cortical areas that mediate attention.

These uses are described in detail in the appendix. One such use is to implement a logical AND operation, such that a particular type of neuron can only be activated if two of its inputs are active, which is supported by compartmental simulations of dendritic spines (Shepherd & Brayton, 1987)

For the sake of simplicity, we assume here that there is a constant level of resistance across the AM, such that voltage is directly proportional to current.

Other models have touched on the idea of simulating the N2pc (Fragopanagos, Kockelkoren, & Taylor 2005) but have not provided a clear link between spatiotopic representations that would produce a lateralized potential

BU and TD were set at .15 and .25 respectively. Higher values of BU allow larger regions to b e selected and vice versa, but the selection of regions is a general property.

This algorithm was chosen for its combination of computational speed, and success in predicting eye-gaze behavior to natural images.

